# Congenital aphantasia is associated with frontotemporal and cingulate structural alterations

**DOI:** 10.64898/2026.03.22.713248

**Authors:** Yusaku Takamura, Romain Delsanti, Laurent Cohen, Paolo Bartolomeo, Jianghao Liu

## Abstract

Congenital aphantasia is characterized by a lifelong self-reported absence of voluntary visual imagery despite preserved visual knowledge, offering a natural model for dissociating sensory representation from conscious imagery. Functional imaging evidence suggests that this dissociation may arise from altered top-down interactions between higher-order control systems and high-level visual cortex, but its structural correlates remain unknown. Here, using diffusion and structural MRI in 18 individuals with congenital aphantasia and 18 matched visualisers, we tested two competing accounts of aphantasia, one predicting structural differences in visual pathways, the other predicting differences in higher-order associative networks. Across multimodal analyses of white-matter tract microstructure, functional-ROI tractography, graph-theoretic network organization, and cortical morphometry, aphantasia was associated with structural differences in frontotemporal and cingulate white-matter tracts, including the uncinate fasciculus, posterior interparietal corpus callosum, and dorsal cingulum, and in frontotemporal cortical regions, including the anterior prefrontal cortex and medial temporal cortex, together forming a candidate neuroanatomical signature for the broad cognitive-affective profile of aphantasia. By contrast, we found no reliable group differences in early visual cortex, major visual tracts, or the direct structural connections of the core imagery network. Congenital aphantasia therefore exhibits a pattern of structural alterations centered on frontotemporal and cingulate systems, with no differences detected in the principal visual structures. Given the modest sample size, these findings are exploratory and hypothesis-generating and require replication in larger cohorts; they nonetheless suggest that higher-order systems supporting integration and conscious access, rather than visual representations themselves, may contribute to the absence of conscious imagery in congenital aphantasia.

## Introduction

Congenital aphantasia, characterized by a lifelong, self-reported absence of voluntary visual mental imagery, affects an estimated 2–4% of the population (Zeman et al., 2015; Zeman, 2024) and presents a striking paradox (Liu & Bartolomeo, 2025; Arcangeli & Bartolomeo, 2025). Aphantasic individuals can accurately retrieve visual properties of objects from memory and perform normally on many visual imagery tasks, yet report little or no subjective imagery experience (Liu & Bartolomeo, 2023; Siena & Simons, 2024; Pounder et al., 2022; Suggate et al., 2026). This dissociation suggests that access to visual knowledge can be preserved even when the phenomenological experience of imagery is absent (Liu & Bartolomeo, 2023). Congenital aphantasia therefore provides a useful model for understanding what higher-order mechanisms are necessary to transform internally generated representations into conscious imagery experience (Liu, 2026).

Beyond the absence of visual imagery, aphantasia appears to reflect heterogeneous, lifelong cognitive and behavioral phenotypes. Functional studies show that aphantasic individuals retrieve fewer phenomenological details during autobiographical recall—a deficit linked to reduced hippocampal–occipital functional connectivity (Dawes et al., 2022; Monzel, Leelaarporn, et al., 2024; Zeman et al., 2020). Beyond memory, impairments are observed in face recognition and emotional awareness (Dance et al., 2023; Monzel, Karneboge, et al., 2024; Zeman et al., 2020), while intrusive memories are significantly less likely to take a visual form (Keogh et al., 2023). Together, these findings indicate that aphantasia may engage distributed, higher-order systems well beyond primary visual representation (Liu, 2026). Consistent with this view, recent frameworks link aphantasia to shared neurodevelopmental mechanisms characterized by widespread reductions in structural and functional connectivity (Sokolowski & Levine, 2023), consistent with co-occurrences with some neurodevelopmental traits and aphantasia (Dance et al., 2021; Mawtus et al., 2024; Milton et al., 2021). However, whether this broad cognitive–affective profile of aphantasia relies on a common anatomical substrate remains unknown. In other neurodevelopmental profiles—such as congenital prosopagnosia, ADHD, and autism spectrum conditions—similar heterogeneous features are underpinned by white-matter microstructural variations in the absence of acquired brain lesions (Parlatini et al., 2023; Thomas et al., 2009; Travers et al., 2012). Structural neuroimaging thus provides a critical means to test whether functional alterations observed in aphantasia share a similar neuroanatomical basis. Specifically, this broad phenotype points to potential structural differences within visual, fronto-temporal and cingulate networks, characterization of which may illuminate both the lifespan stability and phenotypic breadth of aphantasia (Liu, 2026; Zeman et al., 2020).

Neuroimaging studies of aphantasia (Milton et al., 2021; Liu, Cohen, et al., 2025; Chang et al., 2025; Liu, Zhan, et al., 2025; Cabbai et al., 2024; Montabes De La Cruz et al., 2024) have yielded three competing accounts for the absence of imagery experience. First, the sensory strength view holds that aphantasia arises from alterations in early visual areas and their associated visual pathways, reflecting reduced sensory strength of internally generated representations (Keogh & Pearson, 2018). Consistent with this possibility, aphantasic individuals show absence of imagery priming in binocular rivalry (Keogh & Pearson, 2018) and reduction in some physiological signals such as pupil size during imagery tasks (Kay et al., 2022, but see Takamura et al., 2026). In addition, altered cross-decoding during imagery in V1 (Chang et al., 2025) suggests that imagery representations may not reach the perceptual-like format typically associated with vivid imagery experience (Scholz et al., 2025). Second, the conscious access account emphasizes disrupted top-down modulation from higher-order brain regions (Liu, 2026; Liu & Bartolomeo, 2025). During imagery tasks, despite preserved activity in high-level visual areas, aphantasic individuals show reduced functional coupling between the fusiform imagery node (FIN)—a region of fusiform cortex selectively activated during visual imagery (Bartolomeo et al., 2026; Liu, Zhan, et al., 2025; Spagna et al., 2021)—and left prefrontal regions (Liu, Zhan, et al., 2025), including the anterior prefrontal cortex (aPFC) and frontal eye field (FEF). Higher-order attention and salience regions, including FEF and anterior insula, have been proposed to contribute to the top-down amplification of internally generated representations and their segregation from external input (Corbetta & Shulman, 2002). The anterior insula may further gate conscious imagery by regulating activity switching between the dorsal attention and default mode networks (Huang et al., 2021), indicating that conscious access relies on large-scale network dynamics. This account is further supported by reduced representational overlap between imagery and perception in high-level visual regions (Liu, Zhan, et al., 2025) and orbitofrontal cortex (OFC; Liu et al., 2025a), as well as altered frontoparietal-temporal coupling and reduced neural signal complexity (Noble et al., 2026). A third position, the voluntary generation account (Milton et al., 2021; Zeman et al., 2015, 2020), namely, that aphantasia reflects a specific deficit in the voluntary generation of imagery, stems from the observation that many aphantasics report visual dreams despite lacking imagery experience while awake. This account predicts alterations in higher-order brain regions, compatible with the conscious access account, while the latter further provides a mechanistic explanation for how such alterations would abolish imagery experience. The two nonetheless diverge downstream. A deficit in voluntary generation predicts that sensory representations should be absent during attempted imagery, whereas the conscious access account predicts that such representations may be generated but fail to be amplified for awareness. Together, these accounts make different structural predictions. A local visual-pathway account predicts differences in posterior visual tracts, including the vertical occipital fasciculus (VOF), inferior longitudinal fasciculus (ILF), and inferior fronto-occipital fasciculus (IFOF), whereas a higher-order account predicts differences in fronto-temporal and cingulate pathways and their associated large-scale brain networks.

Here, the aim of this study is twofold: first, to identify common neuroanatomical signatures underlying the broad cognitive–affective profile of aphantasia; and second, to test competing predictions regarding the absence of conscious imagery experience. To address both aims, we used multimodal diffusion-weighted and T1-weighted MRI. Specifically, we combined four complementary approaches—tractometry, functional region-of-interest (fROI)-based tractography, graph-theoretic network analysis, and surface-based cortical morphometry—to characterize potential group differences across tract, network, and cortical scales (see analysis pipeline in Fig. S1).

## Results

### Atlas-based tractometry reveals microstructural differences

We quantified fractional anisotropy (FA) along atlas-defined major white-matter tracts, providing a tract-profile measure of white-matter microstructure. We assessed major tracts associated with visual processing, including the VOF, IFOF, and ILF, as well as tracts associated with higher-order functions, including fronto-parietal attention pathways (Superior longitudinal fasciculus: SLFs, Arcuate fasciculus: AF), medial fronto-temporal pathways (Uncinate fasciculus: UF, ventral cingulum), and cognitive-control pathways (dorsal cingulum).

Figure 1 and Table S1 show the tractometry results. No significant FA differences were detected in the visual tracts examined. In contrast, group differences were observed in several higher-order associative pathways. Relative to visualizers, aphantasic individuals showed reduced FA in both the left and right UF, localized to the anterior temporal segment (Fig. 1A). Reduced FA was also observed in the posterior interparietal corpus callosum. By contrast, aphantasic individuals showed increased FA across much of the bilateral dorsal cingulum (Fig. 1B). No significant clusters were identified in the remaining tracts.

**Figure 1.**
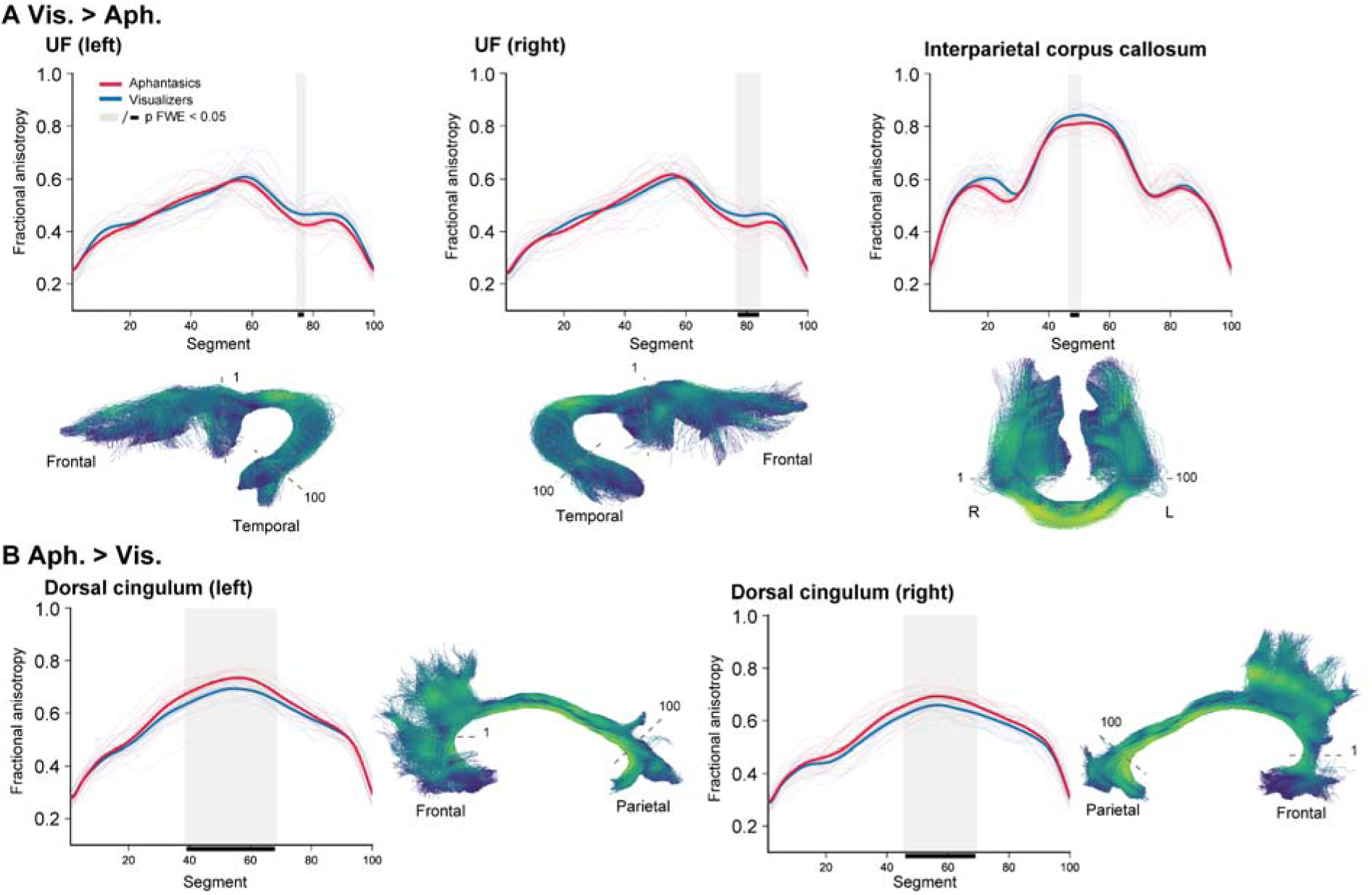
Tractometry reveals white-matter microstructural differences in aphantasia. **(A)** Tracts showing reduced fractional anisotropy (FA) in aphantasic individuals relative to visualizers (Vis. > Aph.): left uncinate fasciculus (UF), right UF, and posterior interparietal corpus callosum. Upper panels show tract profiles (mean ± SEM); lower panels show tract reconstructions from a representative participant, color-coded by segment position (nodes 1-100). Red lines indicate aphantasic individuals and blue lines indicate visualizers. Shaded gray regions mark segments showing significant group differences (p < 0.05, FWE-corrected). **(B)** Tracts showing increased FA in aphantasic individuals relative to visualizers (Aph. > Vis.): left and right dorsal cingulum. Upper panels show tract profiles (mean ± SEM); lower panels show representative tract reconstructions, color-coded by segment position. Red lines indicate aphantasic individuals and blue lines indicate visualizers. Shaded gray regions and black bars on the x axis mark significant segments (p < 0.05, FWE-corrected). Thin lines show individual participant profiles; thick lines show group means.

To further characterize these findings, we examined axial diffusivity (AD), radial diffusivity (RD), and neurite orientation dispersion and density imaging (NODDI) metrics at the affected segments (Fig. S2 and Table S1). These follow-up analyses were consistent with microstructural differences rather than fiber geometry alone, such as crossing-fiber artefact.

The right SLF-II and right ventral cingulum showed marginal reductions in FA (Fig. S3); however, these effects may be confounded by fiber geometry and were therefore not interpreted further.

### Graph-theoretic network analysis reveals disrupted segregation and inter-network hubness

We next applied graph-theoretic analysis to examine whether the tract-level differences extended to the structural organization of large-scale brain networks. We generated a 400 × 400 whole-brain structural connectivity matrix based on the Schaefer 17-network parcellation (see analysi pipeline in Fig. 2A). From this weighted undirected graph, we computed node-level metrics to characterize network organization. Given the study hypothesis, analyses focused on nodes within the visual, attention, salience, and default mode networks.

**Figure 2.**
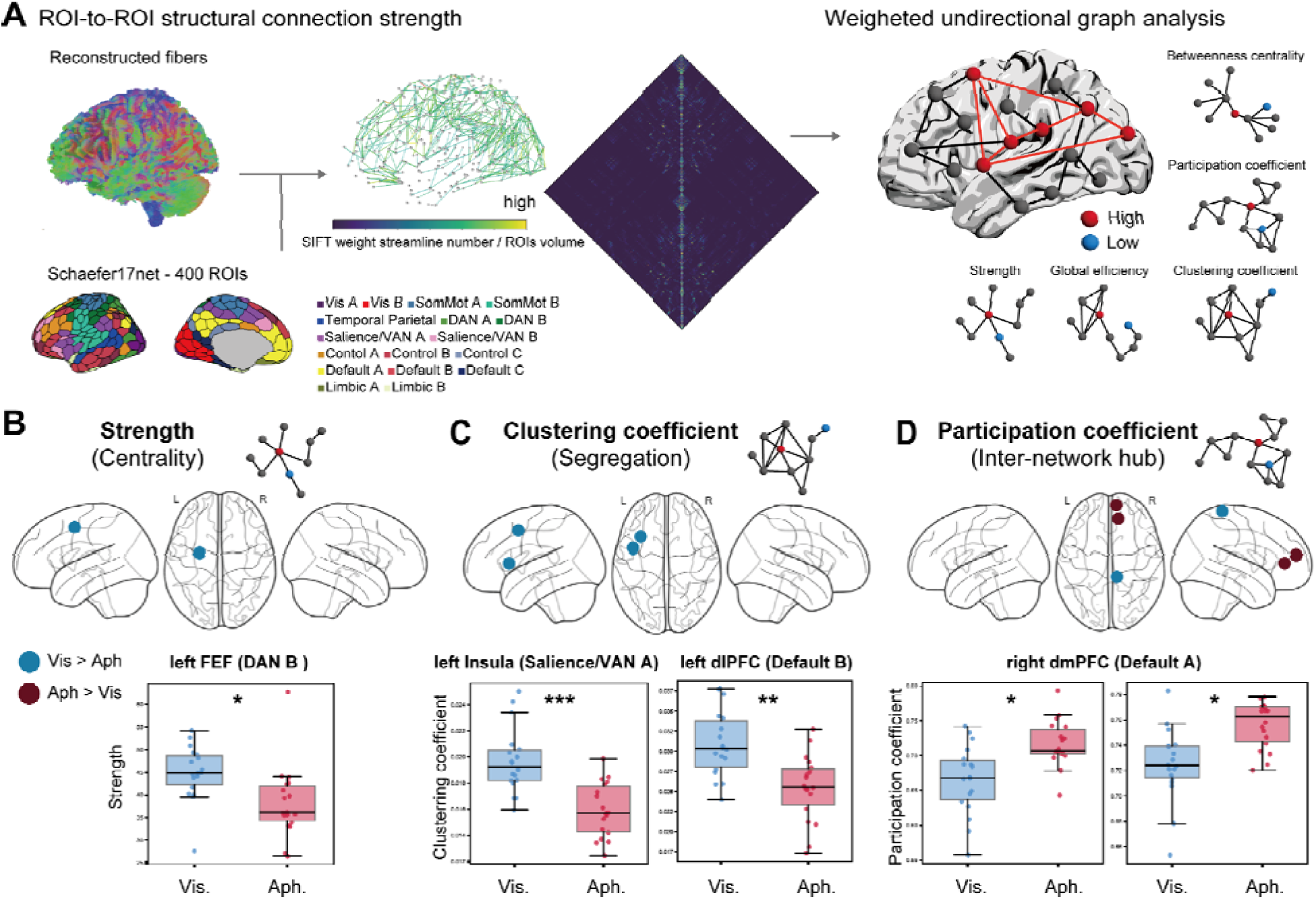
Structural network analysis reveals reduced local segregation and increased inter-network hubness in congenital aphantasia. (A) Analysis pipeline. Whole-brain tractography streamlines were weighted using spherical-deconvolution informed filtering of tractograms (SIFT2) and normalized by ROI volume to generate a 400 × 400 structural connectivity matrix based on the Schaefer 17-network parcellation. Node-level graph-theoretic metrics were then computed from the weighted undirected graph. (B) Node strength was reduced in the left frontal eye field (FEF; dorsal attention network B, DAN B) in aphantasic individuals relative to visualizers, indicating weaker overall structural connectivity at this node. (C) Clustering coefficient was reduced in the left anterior insula (salience/ventral attention network A, Sal/VAN A) and left dorsolateral prefrontal cortex (dlPFC; Default B), indicating reduced local network segregation in salience and association-network regions. (D) Participation coefficient was increased in the right dorsomedial prefrontal cortex (dmPFC; Default A) in aphantasia, indicating greater inter-network hubness at this node. Blue indicates Vis. > Aph.; dark red indicates Aph. > Vis. Boxplots show median and interquartile range; *p < 0.05, **p < 0.01, ***p < 0.001, FWE-corrected.

Group differences were observed in three higher-order network metrics (Fig. 2B–D; Table S2). In aphantasia, the left FEF, a key node of the dorsal attention network (DAN B in Fig. 2A), showed reduced node strength, indicating weaker overall structural connectivity at this node. The left anterior insula, part of the salience/ventral attention network (VAN A), and the left dorsolateral prefrontal cortex (dlPFC) of the default mode network (Default B) showed decreased clustering coefficient, indicating reduced local network segregation in salience and cognitive-control regions. By contrast, the right dorsomedial prefrontal cortex (dmPFC) of the default mode network (Default A) showed increased participation coefficient, indicating stronger inter-network hubness in aphantasic individuals. In addition, the right somatomotor network showed reduced participation coefficient in aphantasic individuals. Control analyses using alternative parcellations confirmed that these findings were robust across atlas options (Fig. S5A–B). Furthermore, a sensitivity analysis on clustering coefficients (see methods) found no significant group differences (Left Insula: B-M statistics = −0.06; Left dlPFC: B-M statistics = 1.30; all pFDR > 0.86), indicating that the insula and dlPFC clustering differences were not specific to network topology. They may reflect differences in the weighted connectome (connection density, strength, or weight distribution) rather than reorganized local clustering per se.

Tracing direct structural connections to these nodes showed that the left anterior insula was connected primarily via the AF and IFOF (Fig. S5C), whereas dmPFC connectivity involved dorsal cingulum pathways. These anatomical relationships are considered further in the Discussion.

By contrast, no significant differences were observed in the visual cortex nodes examined, in keeping with the absence of tractometry differences in the major visual pathways.

### No difference found in white-matter connections of the core imagery network

We next characterized the white-matter connections of a core imagery network whose functional properties are altered in aphantasia. We defined fROIs on the basis of our 7T fMRI results (Liu, Zhan, et al., 2025): the FIN, left aPFC, left OFC, and right anterior intraparietal sulcus (aIPS). For each of the four ROIs (Fig. S4A) in each group, we identified streamlines intersecting its projection on the grey/white-matter interface and quantified tract structure using two indices: volume-normalized streamline count (Fig. S4B) and streamline proportion (Fig. S4C) within anatomically defined bundles. No significant between-group differences were observed for either index in any tract or ROI after correction for multiple comparisons. Bayes factor (BF) analysis further provided some moderate evidence for the absence of a group difference, thus consistent with an equivalence, in volume-normalized streamline count in the VOF connecting the FIN (BF = 0.32), the major white-matter bundle linking dorsal and ventral visual cortex. Together, these findings indicate no reliable alteration in the direct structural connections examined within the core imagery network.

### Cortical thickness: reduced in the left aPFC, increased in limbic regions

We finally examined cortical thickness across the whole brain. Prior work has linked prefrontal and early visual cortical thickness to individual differences in imagery vividness (Bergmann et al., 2016a, 2016b), including a relationship between smaller V1 and stronger imagery (Bergmann et al., 2016b). We therefore examined whether aphantasia is associated with thicker or larger V1.

No significant differences in cortical thickness or volume were observed in any early visual cortex region, with BFs yielding moderate evidence of group equivalence in both the calcarine cortex (Table S3) and V1 (Table S4). By contrast, whole-brain analysis revealed a single significant prefrontal cluster of reduced thickness in the left aPFC in aphantasia (Fig. 3A). Cluster statistics are reported in Table S5. Aphantasic individuals also showed greater cortical thickness in multiple medial temporal limbic regions, including left anterior parahippocampal gyrus and entorhinal cortex; right entorhinal cortex, retrosplenial cortex, posterior cingulate cortex. NODDI analysis identified higher NDI values in bilateral medial temporal regions and the left precuneus in aphantasia (Fig. S6), consistent with microstructural differences in these medial temporal regions.

**Figure 3.**
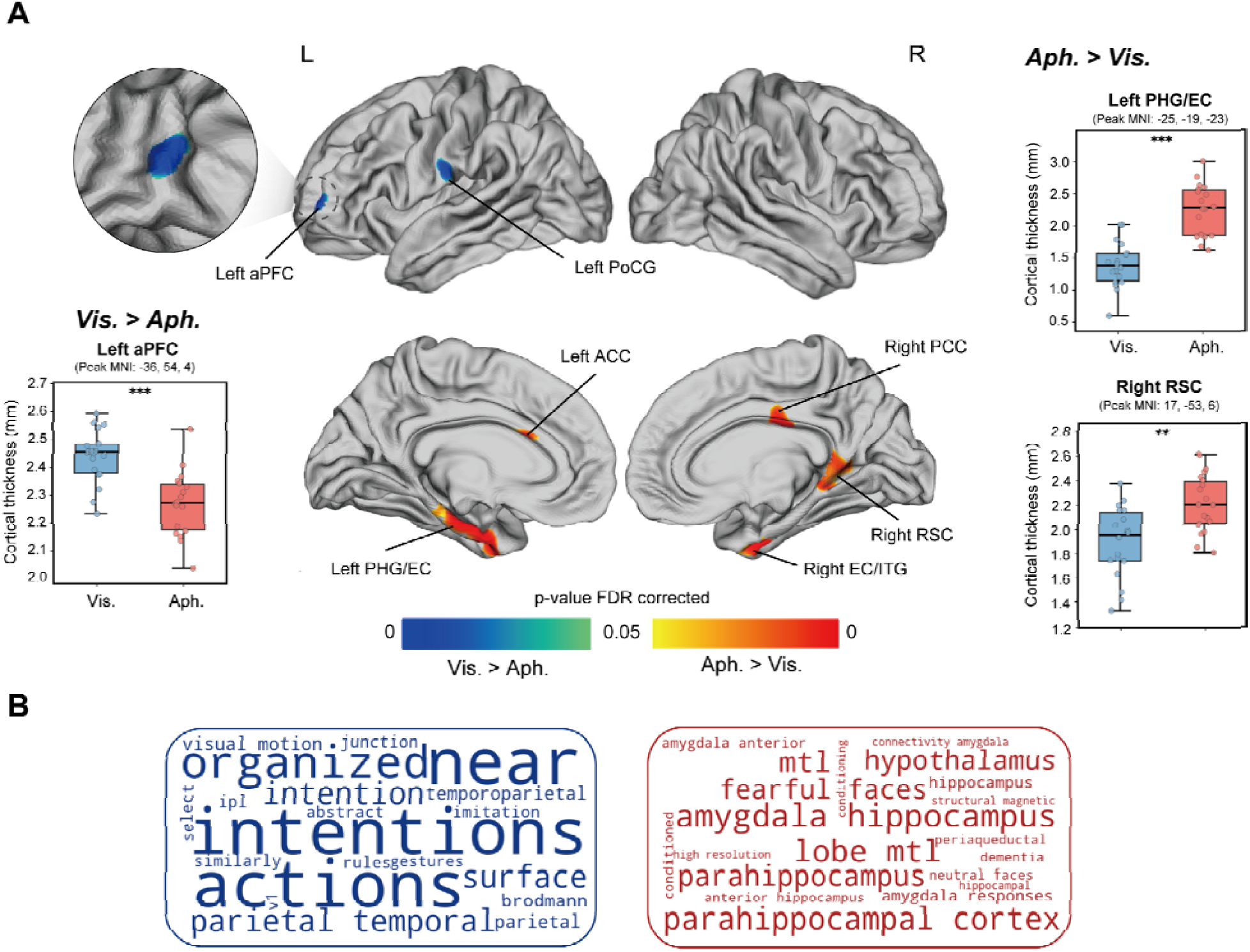
Regional differences in cortical thickness between aphantasic individuals and visualizers. **(A)** Whole-brain map of the group comparison. Blue indicates regions of reduced cortical thickness and red indicates regions of increased cortical thickness in aphantasia, with whole-cortex FDR correction. Boxplots show individual participant values (dots), median, and interquartile range. *p < 0.05, **p < 0.01, ***p < 0.001. Vis., visualizers; Aph., aphantasia. aPFC: anterior prefrontal cortex; PoCG: post central gyrus; PHG: parahippocampal gyrus; EC: entorhinal cortex; ITG: inferior temporal gyrus; RSC: retrosplenial cortex; PCC: posterior cingulate cortex. **(B)** Functional term associations derived from meta-analytic decoding of the t map using Neurosynth. Blue terms are associated with regions of reduced cortical thickness in aphantasia, and red terms with regions of increased cortical thickness. Term size reflects the strength of the Pearson correlation with the corresponding meta-analytic activation map.

To provide functional context for the observed thickness differences, we performed meta-analytic decoding of the t-map using Neurosynth (Fig. 3B). The left aPFC cluster showed strong associations with the terms “intentions” (r = 0.26), “actions” (r = 0.25), and “near (space)” (r = 0.25). Regions of increased thickness in aphantasia showed associations with anatomical terms including hippocampus, parahippocampus, and amygdala (all rs > 0.14), providing convergent functional annotation for the identified regions.

### Multivariate logistic regression

As an exploratory analysis, we used penalized multivariate logistic regression to identify the most parsimonious combinations of structural features associated with group separation. Separate models were fitted for FA, cortical thickness, and graph-theoretical measures, using only features showing group differences in the univariate analyses. The exploratory models showed strong in-sample discrimination, with all areas under the curve (AUCs) > 0.92. Across models, the strongest independent predictors of aphantasia were reduced FA in the right uncinate fasciculus and posterior interparietal corpus callosum, increased FA in the left dorsal cingulum, lower clustering coefficient in the left anterior insula, lower node strength in the left FEF, reduced left aPFC thickness, and greater anterior parahippocampal and retrosplenial cortical thickness. However, the clustering-coefficient difference in the left anterior insula did not survive the null-network sensitivity analysis, so its contribution to the model should be interpreted with caution. Full model statistics are reported in the Supplementary Materials.

Considering the potential heterogeneity of aphantasia, we tested whether the variance of each identified variable differed across groups using Levene’s test for equality of variances, even though all recruited aphantasics reported a lifelong absence of voluntary imagery experience. Variance did not differ significantly between groups (Table S6), although with 18 participants per group this test has limited power to detect heterogeneity. Most participants (13/18) reported core aphantasia, so the present signature is most likely to reflect that subgroup.

## Discussion

Congenital aphantasia has been proposed to reflect reduced functional coupling between higher-order prefrontal systems and high-level visual cortex, rather than a deficit in sensory representation alone (Liu & Bartolomeo, 2025). Here, we report findings that provide structural evidence consistent with this account. Across tractometry, network analysis, and cortical morphometry, aphantasia was associated with a pattern of structural differences centered on fronto-temporal and cingulate systems, whereas we detected no reliable group differences in early visual cortex or in the major visual tracts examined. These findings suggest that congenital aphantasia primarily involves neuroanatomical alterations across distributed but interconnected higher-order brain networks, rather than within a single focal region, potentially underlying its broad cognitive–affective profile.

Four converging observations characterize these structural differences. First, the UF and posterior interparietal corpus callosum showed reduced FA in aphantasia, implicating fronto-temporal and interhemispheric associative pathways. This pattern is consistent with prior functional findings of reduced coupling between OFC and anterior temporal regions in aphantasia (Liu, Cohen, et al., 2025), two territories linked by the UF, potentially reflecting degraded binding between temporal mnemonic content and OFC affective and perceptual processing (Coad et al., 2020; Thomas et al., 2015). It may also be relevant to the broader behavioral profile reported in aphantasia, including altered autobiographical memory, face recognition, and emotional awareness (Zeman et al., 2020; Dawes et al., 2022; Monzel, Leelaarporn, et al., 2024), although direct structure-function relationships remain to be tested. Future studies should therefore assess these structural features alongside objective measures of episodic memory fidelity, face recognition, and affective processing.

Second, increased FA and neurite density in the dorsal cingulum are consistent with altered organization of a cognitive-control pathway. One possibility is that altered top-down regulation affects the conscious accessibility of internally generated representations, in a manner analogous to models of neurofunctional and psychiatric disorders (Mary et al., 2020; Kennis et al., 2015; Sasikumar & Strafella, 2021). Another is that these changes reflect compensatory reliance on semantically guided or non-visual control processes in aphantasia. Distinguishing between these possibilities will require task-based and longitudinal evidence.

Third, at the network level, aphantasia was associated with altered connectivity metrics in frontal association regions, together with increased inter-network hubness in right dmPFC. Reduced segregation was also observed in the left anterior insula and the left dlPFC, but neither difference survived a null-network sensitivity analysis, and both are therefore reported with caution. These findings point to altered organization of attentional, salience, and default-mode systems. Given prior proposals implicating anterior insula and frontal control regions in the top-down regulation of internally generated representations, these network-level differences may affect conscious imagery construction at one or multiple stages (Liu, 2026): mediating inward attentional shifts (Nobre & Gresch, 2025), binding multiple features (Treisman, 2007), and gating conscious access (Huang et al., 2021). Along with the anterior cingulate cortex, insular alterations have also been proposed as a mechanism underlying some cases of acquired aphantasia, via reduced interoceptive and self-referential integration (Scholz et al., 2026; Silvanto & Nagai, 2025).

Fourth, cortical thickness was selectively reduced in the left aPFC, a region which was found to be functionally disconnected in aphantasia (Liu & Bartolomeo, 2025). This anatomical finding, although weaker than the other analyses, provides complementary structural evidence for the involvement of the aPFC in aphantasia and is consistent with its proposed role in higher-order control and conscious access to internally generated representations. The association between aPFC structure and imagery vividness in visualizers (Bergmann et al., 2016b), further supports the relevance of this region to individual differences in imagery. By contrast, cortical thickness was increased in medial temporal limbic regions, including the entorhinal (Moser et al., 2014) and retrosplenial cortex (Vann et al., 2009), critical for spatial navigation, episodic memory and imagined scene construction. Together with the tract and network findings, this pattern localizes the observed structural differences primarily to fronto-temporal and cingulate systems.

Across all four analyses, this structural dissociation is broadly consistent with a recent attention-based multiple-stage model of conscious imagery and aphantasia (Liu, 2026), in which imagery experience depends on at least three broad operations: generation of internal visual representations, integration of distributed visual features into coherent object- or scene-like visual content (see also Scholz et al., 2025; Arcangeli & Bartolomeo, 2025), and amplification of that content for conscious access. Functional neuroimaging converges on the possibility that the first of these processes is at least partly preserved: the visual cortex is reactivated in aphantasic individuals during imagery tasks (Chang et al., 2025; Liu, Cohen, et al., 2025; Liu, Zhan, et al., 2025), consistent with the possibility that sensory reactivation arises from semantic and episodic memory retrieval during imagery generation (Pearson, 2019; Spagna et al., 2024). This interpretation is further supported by our ROI-based analysis, showing similar structural connectivity between the two groups within the core imagery network (Liu, Zhan, et al., 2025). Related behavioural evidence further supports this distinction. During mental exploration of an imagined map of France, aphantasic individuals showed target-directed eye movements similar to visualizers despite reporting little or no imagery vividness (Takamura et al., 2026), suggesting that task-relevant internal representations can be generated even when conscious imagery is limited. Taken together, these findings suggest that generation alone is not sufficient for conscious imagery. Critically, these observations were obtained while participants were deliberately attempting to imagine, and are therefore difficult to reconcile with the prediction of the voluntary generation account (Milton et al., 2021; Zeman et al., 2015, 2020) that sensory representations should be absent under precisely these conditions.

Within this framework (Liu, 2026), altered integration would reduce imagery vividness (see also (Arcangeli & Bartolomeo, 2025; Scholz et al., 2025), whereas altered attentional amplification would gate conscious access through nonlinear processes (Dehaene et al., 2006; Mashour et al., 2020). The aPFC is a plausible candidate for the latter, given its proposed role in perceptual conscious access (Del Cul et al., 2009; Hauw et al., 2024), the reduced functional coupling between the left aPFC and high-level visual cortex reported in aphantasia (Liu & Bartolomeo, 2025), and the reduced cortical thickness of the same region observed in the present study. The involvement of the aPFC might appear to favour the voluntary generation account, given that this region has been implicated in intention and in the attentional control of self-generated cognition (Burgess et al., 2007). We propose, however, that its contribution lies in volitional control and self-reflection over the content of imagery, that is, over which representations are selected and sustained, and with what vividness, rather than in the generation of those representations. Lucid dreaming offers a pertinent dissociation. Dream imagery is produced in the absence of voluntary initiation, yet aPFC engagement during lucid dreaming has been related to the degree of control exerted over the dream and to its vividness and clarity (Baird et al., 2019). A comparable pattern in aphantasia, where visual dreams are reported but are less frequent and qualitatively impoverished (Dawes et al., 2020), is consistent with preserved generation of involuntary imagery alongside reduced aPFC-dependent control over its content and amplification.

On this three-stage account, aphantasia involves relatively preserved sensory generation together with altered integration and conscious access, with attentional amplification potentially representing a more proximal bottleneck for conscious imagery. A recent review noted that, in its present form, this framework does not provide a natural account of the link between aphantasia and impoverished episodic memory (Arnold et al., 2026). The present findings help address this gap, because alterations in the uncinate fasciculus, together with cortical thickness changes in the entorhinal and retrosplenial cortices, implicate mnemonic systems alongside frontotemporal control regions within the broader structural signature of aphantasia. This convergence suggests that alterations affecting mnemonic and frontotemporal control systems may contribute to both imagery and memory phenotypes, providing a potential neuroanatomical link between them. The present structural findings are anatomical, however, and cannot by themselves establish which stage is affected.

The absence of detectable V1 structural differences is consistent with lesion evidence questioning a necessary causal role of the primary visual cortex in visual mental imagery (Bartolomeo et al., 2020). For example, a recent review of 55 cases of visual imagery in cortical blindness reports a double dissociation indicating that visual imagery can be selectively impaired with relative sparing of V1, whereas V1 damage or dysfunction does not necessarily abolish visual imagery (Zago et al., 2026). A recent theoretical proposal (Liu, 2026) holds that V1 activity is shaped by feedback-driven inhibition of spontaneous activity (see also Koenig-Robert et al., 2026), selectively suppressing signals that do not encode imagined content. A balanced excitatory-inhibitory regime in V1 would then serve to prevent interference from external input and sustain attention inward, stabilising internally generated representations. On this view, the altered V1 activity reported in aphantasia (see Liu, 2026, for review) would reflect a functional imbalance arising from altered modulation by higher-order networks rather than a structural deficit within V1 itself. In the present data, we did not detect group differences in these regions, and Bayes factors favoured equivalence in several, though not all, of the early visual subregions and tracts examined. This does not fully exclude visual-pathway involvement. The visual-pathway account remains open and requires more sensitive investigation, including of superficial U-shaped fibres, before it can be evaluated.

A notable feature is that the direction of the neuroanatomical alterations was not uniform across measures and therefore cannot be interpreted as a single pattern of structural deficit. Interestingly, neuroanatomical alterations in the direction of increase in aphantasia were predominantly localized to medial cortical and cingulate territories, including the dorsal cingulum, cingulate cortex, and medial prefrontal cortex. This spatial distribution may indicate altered organization of medial control and mnemonic systems involved in cognitive control and executive functions (Bubb et al., 2018; Petersen & Posner, 2012). Whether these alterations reflect a functionally advantageous or compensatory form of neuroanatomical reorganization remains unclear. A previous study found that properties of the dorsal cingulum were positively associated with response inhibition in pediatric obsessive-compulsive disorder, but not in healthy controls, raising the possibility that alterations in this pathway may, under some conditions, reflect compensatory reorganization (Gruner et al., 2012). Consistent with this possibility, individuals with aphantasia have also been reported to employ alternative strategies during tasks that typically require visual imagery (Reeder et al., 2024), and to show increased resting state functional connectivity between the anterior cingulate cortex and hippocampus (Milton et al., 2021). Thus, both decreases and increases in neuroanatomical signatures may reflect distinct aspects of the neural organization underlying aphantasia.

Importantly, the cognitive interpretation of these neuroanatomical findings remains indirect. Although the observed alterations involved frontotemporal, cingulate, salience, and attentional networks implicated in higher-order control, internal monitoring, and conscious access, these functions were not independently assessed in the same participants. Future studies combining neuroanatomical measures with direct cognitive and behavioural assessments will be important for linking these alterations to individual differences in imagery and associated cognitive functions.

Some limitations should be noted. First, given the substantial phenotypic heterogeneity of aphantasia (Delem et al., 2025; Nanay, 2025), larger and more extensively phenotyped cohorts will be needed to assess the generalizability of the present findings and to identify features shared across heterogeneous phenotypes. The variance did not differ between groups, and most participants reported core aphantasia, so the neuroanatomical signature reported here may capture a central characteristic of the condition. Second, although our multimodal analyses provide convergent evidence for frontotemporal and cingulate involvement, the tract-profile microstructure and graph-theoretical analyses share the same diffusion MRI acquisition and preprocessing pipeline, and diffusion MRI-derived measures provide model-based estimates rather than direct measures of their underlying biological substrates (F. Zhang et al., 2022). The cortical thickness findings, by contrast, were derived from an independent T1-weighted modality. Third, the cortical thickness on left aPFC is identified in a whole-brain analysis and is substantially weaker than the other analysis streams. Its interpretive value lies in convergence with independent functional evidence implicating the same region (Liu & Bartolomeo, 2025) rather than standing alone.

In summary, this exploratory study provides a multimodal characterization of structural brain differences in congenital aphantasia, centred on higher-order fronto-temporal and cingulate systems - a candidate neuroanatomical signature for the broad cognitive-affective profile of the condition. By contrast, we found little evidence for major structural differences in early visual cortex or in its principal white-matter tracts, although the status of local U-shaped fibers (Guevara et al., 2020) remains to be determined. These findings point to a structural basis of conscious imagery that might lie not in sensory representations themselves, but in the higher-order integrative and control systems that enable access to them, although the present data cannot exclude a contribution of visual pathways.

## Contributions

**Jianghao Liu**: Conceptualization, Data curation, Formal analysis, Funding acquisition, Investigation, Methodology, Project administration, Supervision, Visualization, Writing – original draft, Writing – review & editing; **Yusaku Takamura**: Conceptualization, Data curation, Formal analysis, Methodology, Software, Visualization, Writing – original draft, Writing – review & editing; **Paolo Bartolomeo:** Conceptualization, Funding acquisition, Project administration, Resources, Supervision, Writing – review & editing. **Laurent Cohen:** Resources, Writing – review & editing; **Romain Delsanti**: Data curation, Writing – review & editing.

## Code and data availability

All derived data and analysis code required to reproduce the results are publicly available at Zenodo under a CC-BY 4.0 licence: https://doi.org/10.5281/zenodo.22798825. Raw and preprocessed MRI data are not publicly shared due to data protection regulation. For any further request and correspondence, please contact the corresponding author.

## Acknowledgement

We gratefully thank all CENIR collaborators for their technical assistance in data acquisition and for access to the neuroimaging platform. We thank all the members of the PICNIC lab for helpful discussion and suggestions. We thank Dounia Hajhajate, Hanna Kavaliova, and Diane Lamarle for their help in collecting part of the data, Minye Zhan, Elena Grosso, and Benoît Béranger for their suggestions regarding the data analysis and Michel Thiebaut de Schotten for commenting on an early version of our results. The work of Jianghao Liu is supported by specific funding from Dassault Systèmes. The work of Yusaku Takamura is supported by JSPS KAKENHI Grants JP24KK0296 and JST-CREST JPMJCR23P1. The work of Paolo Bartolomeo is supported by the Agence Nationale de la Recherche through ANR-16-CE37-0005 and ANR-10-IAIHU-06, by the Fondation pour la Recherche sur les AVC through FR-AVC-017, and by the Paris Brain Institute grant ViBER - Vision Beyond External Reality.

## Supplementary materials

**Supplementary Figure S1.**
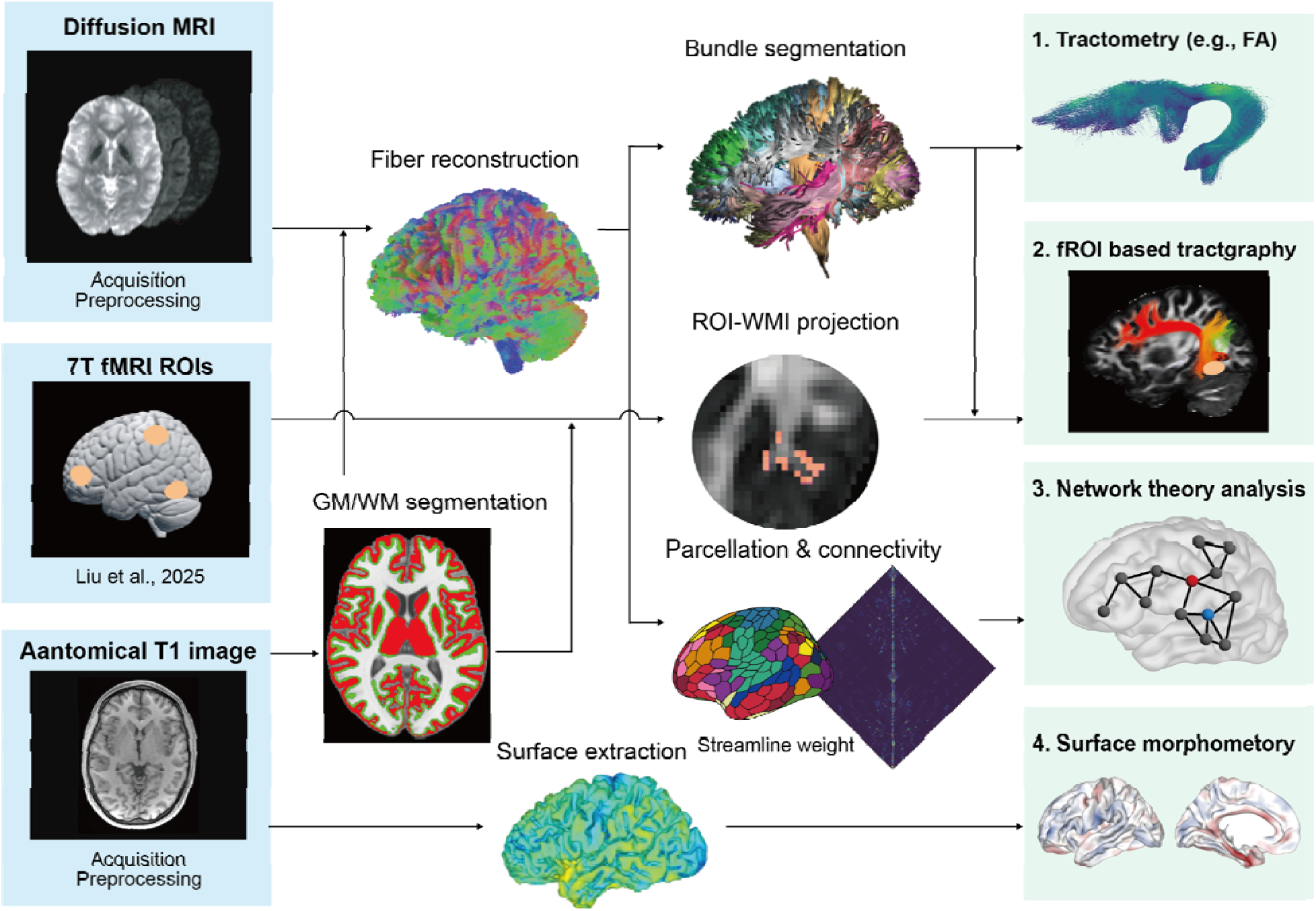
Overview of the analysis pipeline. Three data streams were processed in parallel. Diffusion MRI data underwent preprocessing and fiber reconstruction to generate whole-brain tractography. Anatomical T1-weighted images were processed for grey matter/white matter (GM/WM) segmentation and cortical surface extraction. Functional ROIs from a previous 7T fMRI study of mental imagery (Liu, Zhan, et al., 2025) were registered to each participant’s native space. These inputs were then used for four complementary analyses: **(1) tractometry**, in which white-matter bundles were segmented and microstructural properties, including fractional anisotropy (FA), were quantified along tract profiles; **(2) fROI-based tractography**, in which functional ROIs were projected to the GM/WM interface to identify streamlines connecting regions of the core imagery network; **(3) graph-theoretic network analysis**, in which cortical parcellation and SIFT2-weighted streamline counts were used to construct structural connectivity matrices and derive node-level network metrics; and **(4) surface morphometry**, in which cortical thickness was estimated from the extracted cortical surfaces and compared between groups across the whole brain.

### Axial and radial diffusivity and NODDI control analyses for tractometry findings

To clarify the potential biological basis of the FA differences, we examined axial diffusivity (AD) and radial diffusivity (RD), often interpreted as being more sensitive to axonal and myelin-related properties, respectively (Song et al., 2002), together with neurite orientation dispersion and density imaging (NODDI) metrics, including neurite density index (NDI) and orientation dispersion index (ODI) (H. Zhang et al., 2012), at all segments showing significant FA effects. Full statistics are reported in Table S1.

#### Uncinate fasciculus and posterior interparietal corpus callosum

FA reductions in the bilateral uncinate fasciculus and posterior interparietal corpus callosum were accompanied by marginal RD increases at largely overlapping segments (all p < 0.1, FWE-corrected), a pattern consistent with altered myelin-related microstructure. No significant group differences in NDI or ODI were observed at the same segments, suggesting that these FA reductions are unlikely to be explained by fiber-geometry effects such as crossing fibers.

#### Bilateral dorsal cingulum

The FA increase in the bilateral dorsal cingulum was accompanied, in the left hemisphere, by a significant RD decrease and AD increase, consistent with altered microstructural organization rather than myelin degradation. A significant increase in NDI was observed at the same segments showing elevated FA, with no change in ODI, suggesting that the FA increase is more likely to reflect greater neurite density than reduced fiber dispersion.

#### Marginal findings

Marginal FA reductions were observed in the right SLF-II (Fig. S3A; p = 0.082) and right ventral cingulum (p = 0.079). Both tracts also showed increased ODI at the same segments (Fig. S3B), suggesting that these marginal effects may be influenced by local fiber geometry rather than robust microstructural differences. We therefore did not interpret these findings further.

**Supplementary Figure S2.**
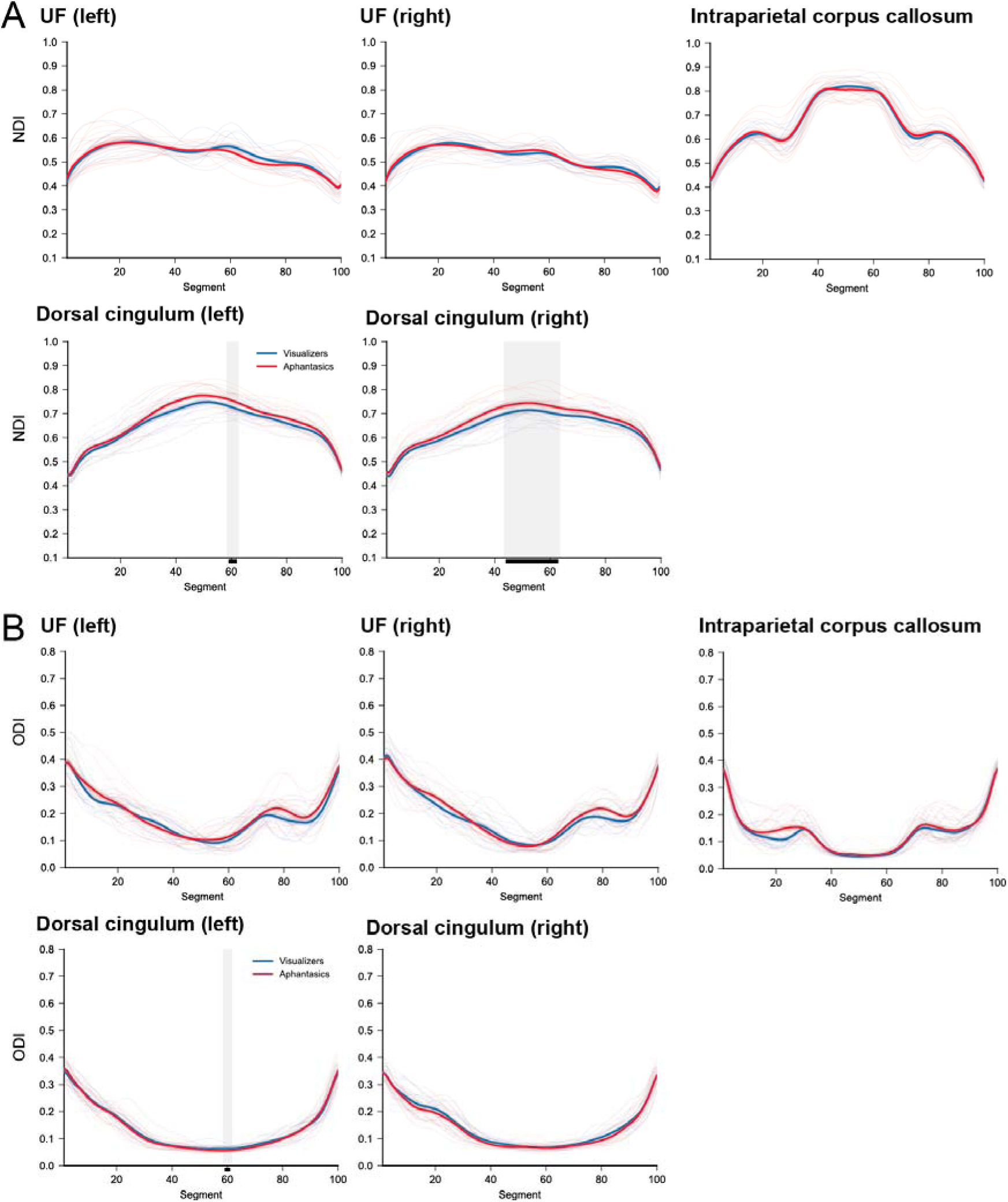
NODDI microstructural profiles in white-matter tracts showing significant FA differences. **(A)** Neurite density index (NDI) tract profiles. **(B)** Orientation dispersion index (ODI) tract profiles. Profiles are shown for the bilateral uncinate fasciculus, posterior interparietal corpus callosum, and bilateral dorsal cingulum, the tracts identified in the main tractometry analysis. Red lines indicate aphantasic individuals and blue lines indicate visualizers. Thin lines show individual participant profiles; thick lines show group means ± SEM. Shaded grey regions and black bars indicate significant between-group differences (p < 0.05, FWE-corrected).

**Supplementary Figure S3.**
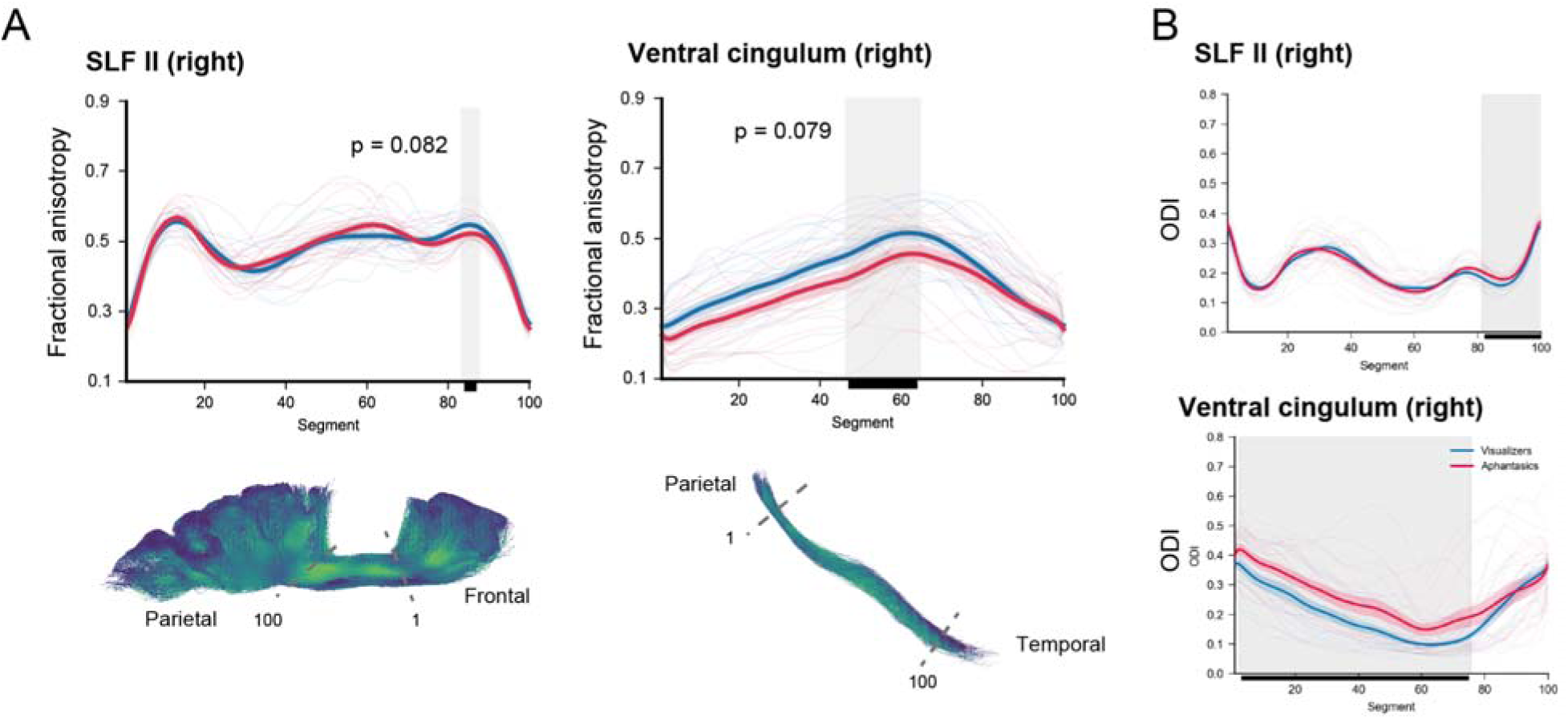
Marginal FA reductions in the right SLF-II and right ventral cingulum are accompanied by increased orientation dispersion. **(A)** Fractional anisotropy (FA) tract profiles for the right SLF-II (p = 0.082, FWE-corrected) and right ventral cingulum (p = 0.079, FWE-corrected), with representative tract reconstructions shown below. Shaded grey regions and black bars indicate segments showing marginal between-group differences. Red line indicate aphantasic individuals and blue lines indicate visualizers; thin lines show individual participant profiles and thick lines show group means ± SEM. **(B)** Orientation dispersion index (ODI) profiles for the same tracts. In both tracts, marginal FA reductions overlapped with increased ODI in aphantasic individuals, suggesting that these effects may be influenced by local fiber geometry rather than robust microstructural differences. These findings were therefore not interpreted further.

**Supplementary Figure S4.**
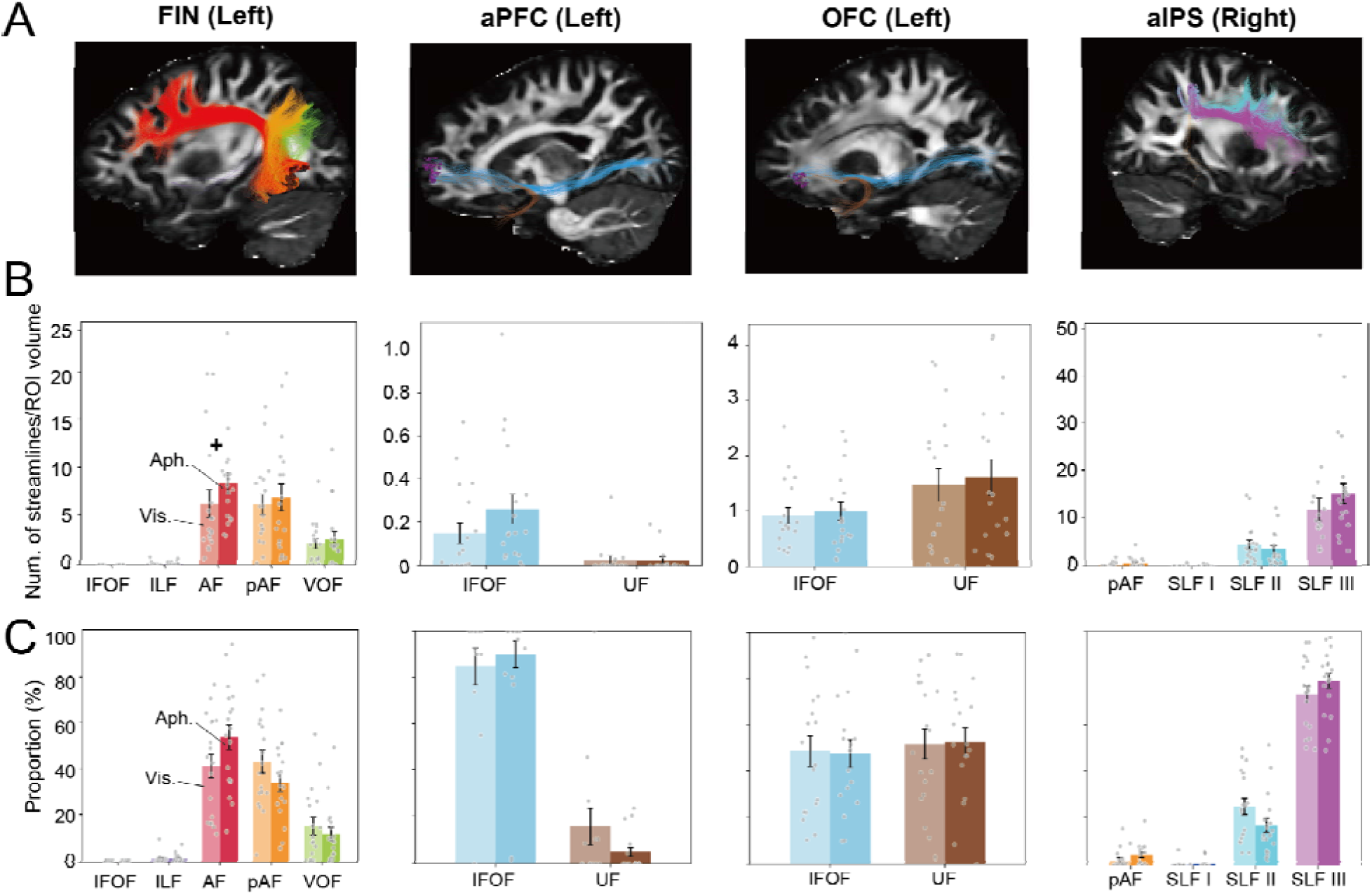
White-matter connections of the core imagery network. **(A)** White-matter streamlines associated with each functional ROI of the core imagery network, shown on sagittal MRI sections: left fusiform imagery node (FIN), left orbitofrontal cortex (OFC), left anterior prefrontal cortex (aPFC), and right anterior intraparietal sulcus (aIPS). **(B)** Streamline counts normalized by ROI volume for each functional ROI. The FIN was connected primarily by the long segment of the arcuate fasciculus (AF), posterior arcuate fasciculus (pAF), and vertical occipital fasciculus (VOF). The left OFC was connected by both the inferior fronto-occipital fasciculus (IFOF) and uncinate fasciculus (UF), whereas the left aPFC was connected mainly by the IFOF. No direct structural connection was identified between the FIN and left aPFC in this analysis. The right aIPS was connected predominantly by the third branch of the superior longitudinal fasciculus (SLF-III). A nominal increase in normalized streamline count was observed in aphantasic individuals for the left AF associated with the FIN (uncorrected p = 0.039, U = 229, Cliff’s d = −0.41), but this effect did not survive correction for multiple comparisons. Bayes factor analysis provided moderate evidence for no group difference in VOF streamline count associated with the FIN (BF = 0.32). **(C)** Proportion of streamlines assigned to each tract of interest for each ROI. Light bars indicate visualizers and dark bars indicat aphantasic individuals. Error bars show SEM and dots indicate individual participants. IFOF, inferior fronto-occipital fasciculus; ILF, inferior longitudinal fasciculus; AF, arcuate fasciculus long segment; pAF, posterior arcuate fasciculus; VOF, vertical occipital fasciculus; UF, uncinate fasciculus; SLF-I/II/III, superior longitudinal fasciculus branches I, II, and III.

### Control analysis on atlas selection in network analysis

**Supplementary Figure S5.**
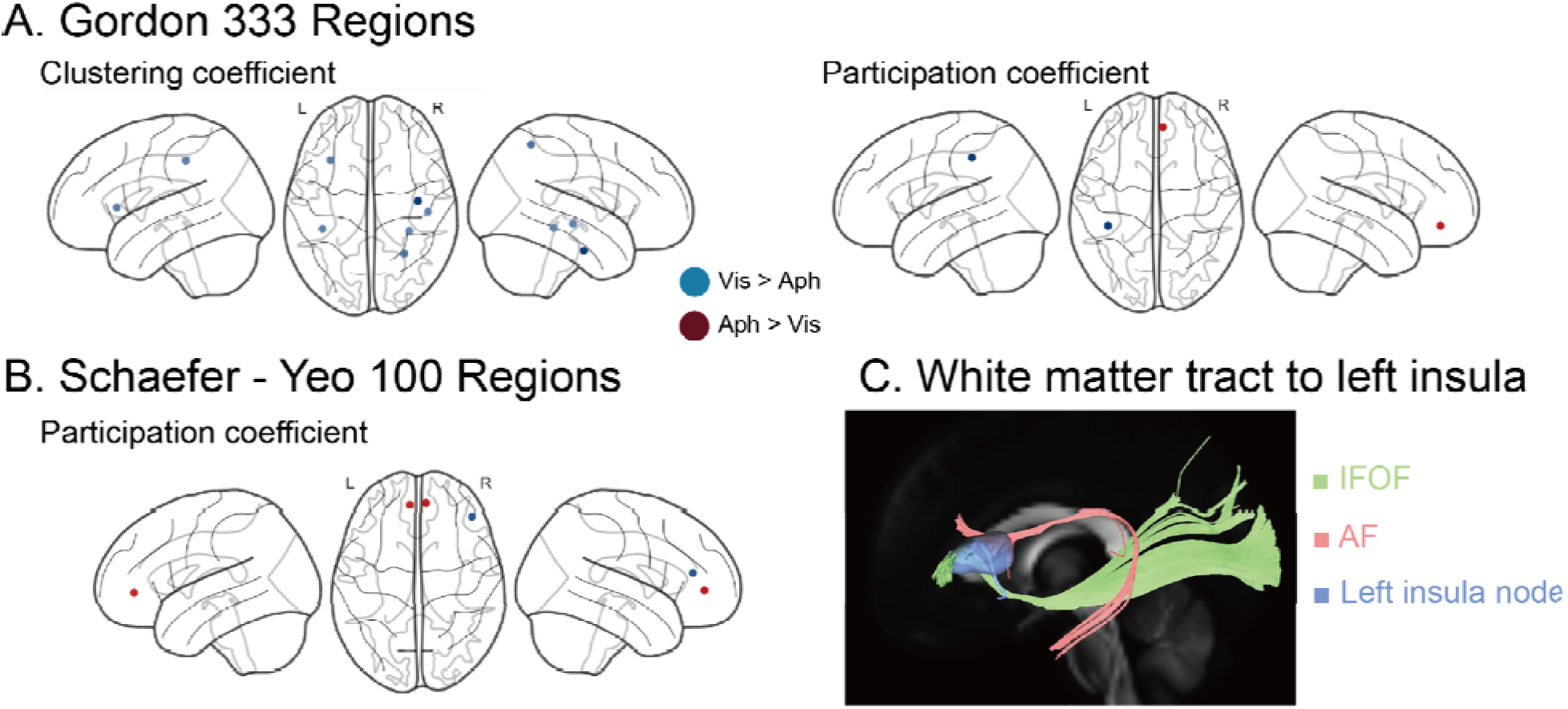
Control analyses confirm the robustness of network findings across parcellation schemes. **(A)** Gordon 333-region atlas. Convergent group differences were observed across graph-theoretic metrics, with aphantasic individuals showing reduced clustering coefficient in the left anterior insula (Vis. > Aph., blue) and increased participation coefficient in the right dorsomedial prefrontal cortex (dmPFC; Aph. > Vis., dark red). **(B)** Schaefer-Yeo 100-region atlas. Aphantasic individuals showed increased participation coefficient in bilateral dmPFC (Aph. > Vis.) and reduced participation coefficient in right lateral prefrontal cortex (Vis. > Aph.). **(C)** White-matter tracts connected to the left anterior insula node in the Schaefer-Yeo 400-region atlas. The inferior fronto-occipital fasciculus (IFOF) and arcuate fasciculus (AF) converge on the left anterior insula, the node showing reduced clustering coefficient in the main analysis. The left anterior prefrontal cortex (aPFC) and fusiform imagery node (FIN) are linked by distinct white-matter pathways, primarily IFOF and AF, respectively. Dark red indicates Aph. > Vis. and blue indicates Vis. > Aph. All group differences shown in panels A and B were significant at p < 0.05, FDR-corrected. The left frontal eye field (FEF) was not identified consistently across atlases, consistent with uncertainty in its parcellation boundaries.

### Control analysis on NODDI gray matter

**Supplementary Figure S6.**
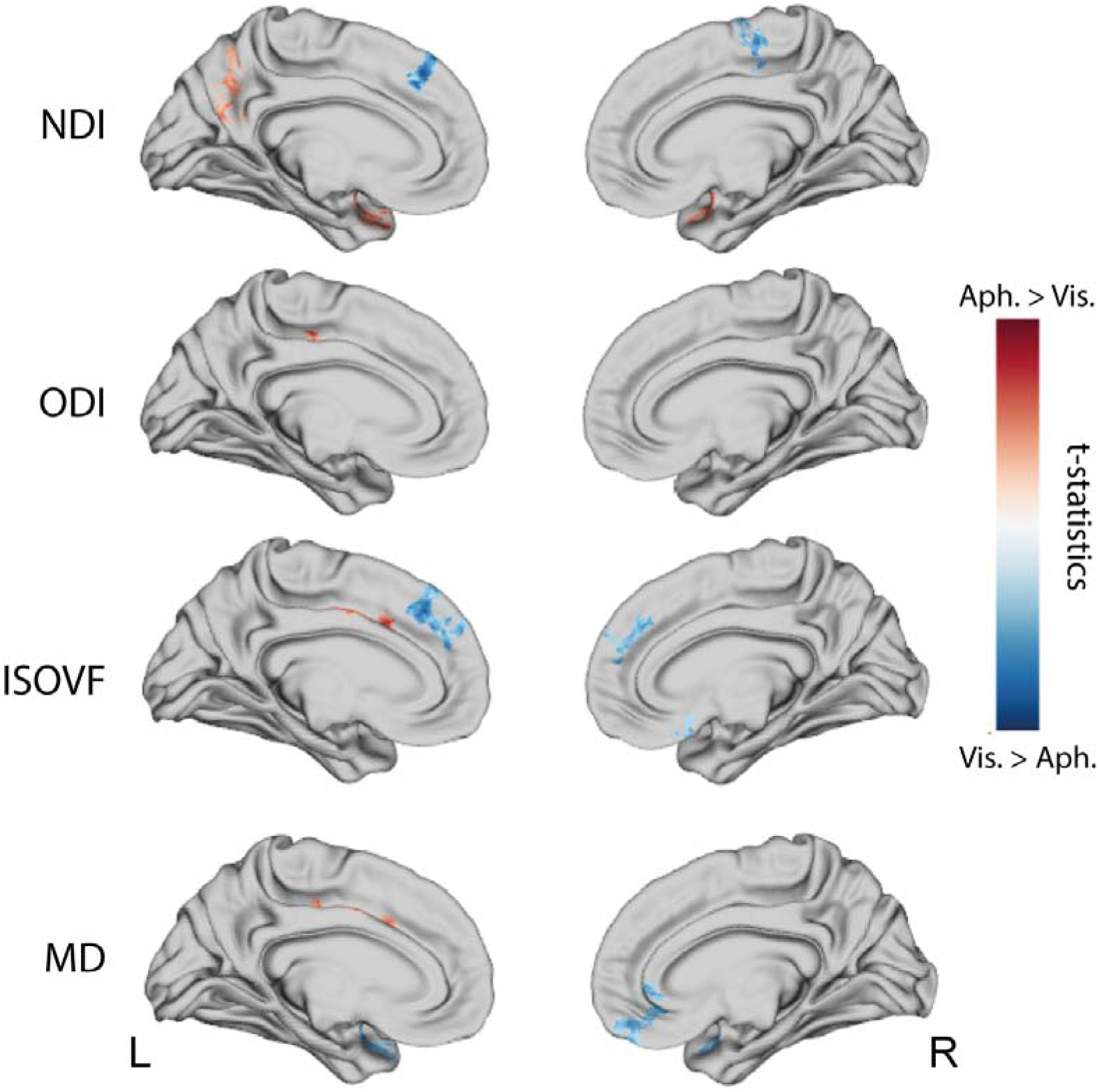
Cortical NODDI microstructural differences between aphantasic individuals and visualizers. Cortical NODDI metrics were compared between groups using random field theory as implemented in BrainStat, with cluster-level FWE correction (p < 0.05). Aphantasic individuals showed increased neurite density index (NDI) in bilateral medial temporal regions, including the parahippocampal gyrus and entorhinal cortex, as well as in the left precuneus, and reduced NDI in the left dorsomedial prefrontal cortex. Orientation dispersion index (ODI), isotropic volume fraction (ISOVF), and mean diffusivity (MD) also showed significant group differences. Regions of reduced MD overlapped spatially with areas of increased NDI in medial temporal cortex, a pattern consistent with denser neurite packing and lower overall diffusivity in these regions.

**Supplementary Table S1.**
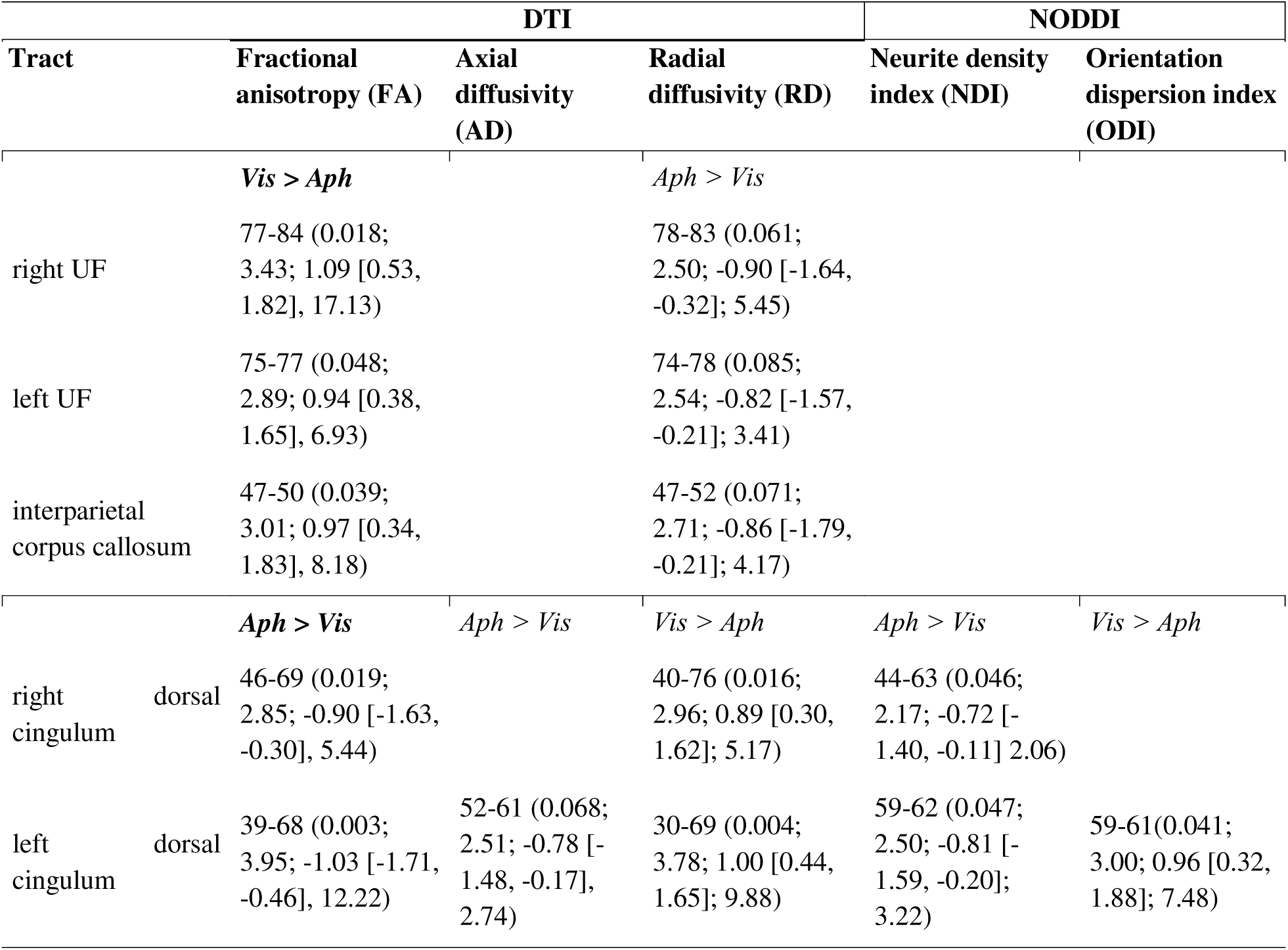
Tract-level diffusion and NODDI differences between aphantasic individuals and visualizers. Significant group differences in tract-profile diffusion tensor imaging (DTI) and neurite orientation dispersion and density imaging (NODDI) metrics are shown for tracts identified in the tractometry analysis. Values indicate the start and end nodes of each significant cluster, followed by the FWE-corrected p value, test statistic, Hedges’ g with bootstrap 95% confidence interval, and Bayes Factor in parentheses. Direction of effect is indicated separately for each metric column (Vis. > Aph. or Aph. > Vis.). Only clusters meeting the reporting threshold of at least 3 consecutive significant nodes are listed. For the main analyses, significance was defined as p < 0.05, FWE-corrected; marginal effects (p < 0.1, FWE-corrected) are reported in the text but are not included in this table.

| Tract | DTI |  |  | NODDI |  |
|---|---|---|---|---|---|
| Tract | Fractional anisotropy (FA) | Axial diffusivity (AD) | Radial diffusivity (RD) | Neurite density index (NDI) | Orientation dispersion index (ODI) |
|  | ***Vis > Aph*** |  | *Aph > Vis* |  |  |
| right UF | 77-84 (0.018; 3.43; 1.09 [0.53, 1.82], 17.13) |  | 78-83 (0.061; 2.50; -0.90 [-1.64, -0.32]; 5.45) |  |  |
| left UF | 75-77 (0.048; 2.89; 0.94 [0.38, 1.65], 6.93) |  | 74-78 (0.085; 2.54; -0.82 [-1.57, -0.21]; 3.41) |  |  |
| interparietal corpus callosum | 47-50 (0.039; 3.01; 0.97 [0.34, 1.83], 8.18) |  | 47-52 (0.071; 2.71; -0.86 [-1.79, -0.21]; 4.17) |  |  |
|  | ***Aph > Vis*** | *Aph > Vis* | *Vis > Aph* | *Aph > Vis* | *Vis > Aph* |
| right dorsal cingulum | 46-69 (0.019; 2.85; -0.90 [-1.63, -0.30], 5.44) |  | 40-76 (0.016; 2.96; 0.89 [0.30, 1.62]; 5.17) | 44-63 (0.046; 2.17; -0.72 [-1.40, -0.11] 2.06) |  |
| left dorsal cingulum | 39-68 (0.003; 3.95; -1.03 [-1.71, -0.46], 12.22) | 52-61 (0.068; 2.51; -0.78 [-1.48, -0.17], 2.74) | 30-69 (0.004; 3.78; 1.00 [0.44, 1.65]; 9.88) | 59-62 (0.047; 2.50; -0.81 [-1.59, -0.20]; 3.22) | 59-61(0.041; 3.00; 0.96 [0.32, 1.88]; 7.48) |

**Supplementary Table S2.** Nodal graph-theoretic network differences between aphantasic individuals and visualizers. Significant group differences in nodal graph-theoretic metrics are reported for node strength, clustering coefficient, and participation coefficient. Group comparisons were performed using the Brunner-Munzel test with false discovery rate (FDR) correction within each metric. MNI coordinates (x, y, z) correspond to parcel centroids. Network affiliations are defined according to the Schaefer 400-parcel, 17-network atlas. **Vis. > Aph.** indicates lower metric values in aphantasic individuals, whereas **Aph. > Vis.** indicates higher metric values in aphantasic individuals. **B-M statistic** denotes the Brunner-Munzel statistic, and **Cliff’s delta** the effect size. Significance threshold: p < 0.05, FDR-corrected.

| Metric | Node (Network) | MNI coordinates (x, y, z) | B-M statistic | p(FDR corrected) | Cliff’s delta [Bootstrap 95%CI] | BF10 |
|---|---|---|---|---|---|---|
| ***Vis. > Aph.*** |  |  |  |  |  |  |
| Strength | Left FEF (DAN B) | -25, -1, 55 | -4.33 | 0.050 | 0.65<br>[0.33, 0.92] | 8.24 |
| Clustering coefficient | Left anterior insula (Salience/VAN A) | -33, 18, 6 | -7.18 | <0.001 | 0.77<br>[0.53, 0.94] | 72.90 |
| Clustering coefficient | Left dlPFC (Default B) | -42, 7, 48 | -4.93 | 0.004 | 0.67<br>[0.37, 0.90] | 19.96 |
| Participation coefficient | Right somatomotor (SomMot B) | 9, -40, 68 | -4.05 | 0.038 | 0.60<br>[0.28, 0.85] | 8.71 |
| ***Aph. > Vis.*** |  |  |  |  |  |  |
| Participation coefficient | Right dmPFC (Default A) | 7, 42, 4 | 4.64 | 0.024 | -0.67<br>[-0.91, -0.36] | 9.68 |
| Participation coefficient | Right dmPFC (Default A) | 7, 54, 13 | 4.19 | 0.038 | -0.63<br>[-0.89, -0.31] | 15.88 |

**Supplementary Table S3.** Gray matter volume in occipital cortical regions. Regions were defined using the Neuromorphometrics atlas in CAT12. Gray matter volumes were normalized by total intracranial volume (TIV). No significant group differences were observed in any occipital cortical region. Bayes factors provided moderate evidence for an equal volume between the two groups in the left and right calcarine cortices. Values are reported as mean ± SD.

| **Region** | **Aphantasics<br>(× 10^-4 mm<sup>3</sup>/TIV)** | **Visualizers<br>(× 10^-4 mm<sup>3</sup>/TIV)** | **BF+0** |
|---|---|---|---|
| Right calcarine cortex | 20 ± 2.26 | 20 ± 2.70 | **0.21\*** |
| Left calcarine cortex | 20 ± 2.71 | 20 ± 2.35 | **0.22\*** |
| Right inferior occipital gyrus | 40 ± 2.50 | 40 ± 3.15 | 0.35 |
| Left inferior occipital gyrus | 30 ± 2.97 | 30 ± 3.74 | 0.33 |
| Right lingual gyrus | 50 ± 4.74 | 50 ± 3.72 | 0.75 |
| Left lingual gyrus | 50 ± 2.87 | 50 ± 3.65 | 0.59 |
| Right middle occipital gyrus | 30 ± 3.50 | 30 ± 2.84 | **0.23\*** |
| Left middle occipital gyrus | 40 ± 4.50 | 40 ± 2.94 | **0.33\*** |
| Right occipital pole | 20 ± 2.94 | 20 ± 2.67 | 0.42 |
| Left occipital pole | 20 ± 2.18 | 20 ± 2.21 | **0.28\*** |
| Right occipital fusiform gyrus | 20 ± 2.10 | 20 ± 2.13 | 0.92 |
| Left occipital fusiform gyrus | 20 ± 1.89 | 20 ± 2.10 | 0.37 |

**Note:** For all Bayesian tests, the alternative hypothesis specified smaller regional volume in visualizers than in aphantasics. **\*BF<sub>+0</sub> < 0.33**, indicating moderate evidence for the null hypothesis.

**Supplementary Table S4.**
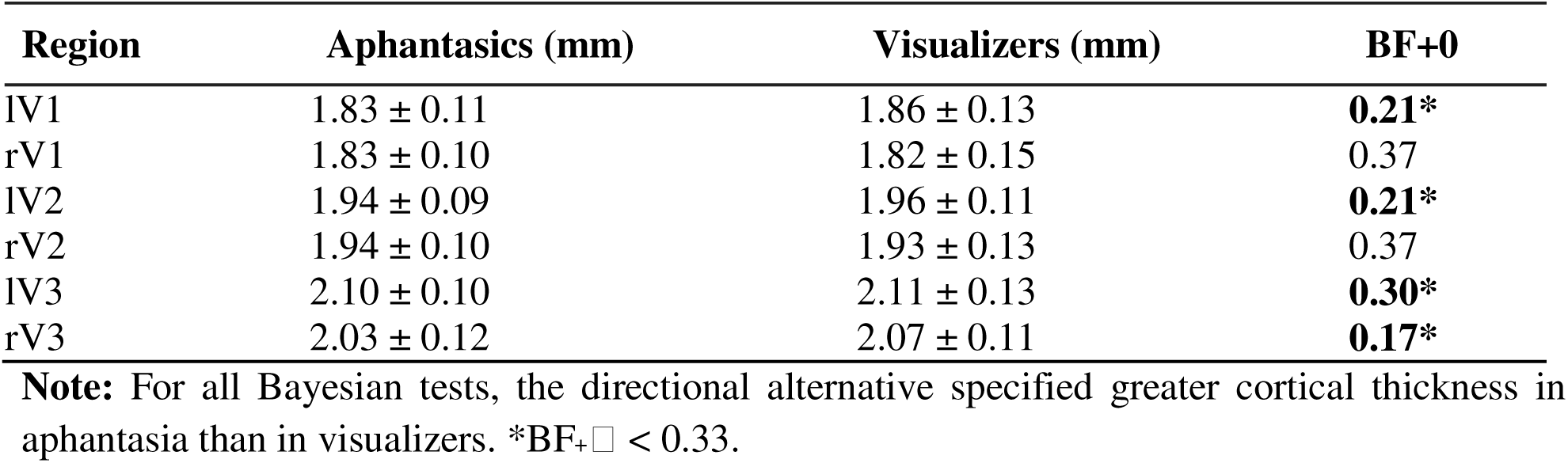
Cortical thickness in early visual areas in aphantasic individuals and visualizers. Mean cortical thickness values (mm ± SD) for atlas-defined early visual regions are shown for the aphantasia and visualizer groups. Regions of interest were defined using the HCP-MMP atlas. No significant group differences were observed in any region. Bayes factors indicated moderate evidence for the absence of a group difference in 4 of the 6 regions (BF_+_ < 0.33). l, left hemisphere; r, right hemisphere.

| Region | Aphantasics (mm) | Visualizers (mm) | BF+0 |
|---|---|---|---|
| lV1 | 1.83 ± 0.11 | 1.86 ± 0.13 | **0.21\*** |
| rV1 | 1.83 ± 0.10 | 1.82 ± 0.15 | 0.37 |
| lV2 | 1.94 ± 0.09 | 1.96 ± 0.11 | **0.21\*** |
| rV2 | 1.94 ± 0.10 | 1.93 ± 0.13 | 0.37 |
| lV3 | 2.10 ± 0.10 | 2.11 ± 0.13 | **0.30\*** |
| rV3 | 2.03 ± 0.12 | 2.07 ± 0.11 | **0.17\*** |

**Note:** For all Bayesian tests, the directional alternative specified greater cortical thickness in aphantasia than in visualizers. \*BF<sub>+</sub>□ < 0.33.

**Supplementary Table S5.** Cortical thickness differences between aphantasic individuals and visualizers. Significant group differences in cortical thickness are reported for clusters comprising more than 30 vertices. Group comparisons were performed using a linear model with FDR correction. aPFC: anterior prefrontal cortex; PoCG: Post central gyrus; PHG: parahippocampal gyrus; EC: Entorhinal cortex; ACC: Anterior cingulate cortex; RSC: retrosplenial cortex; PCC: Posterior cingulate cortex.

| **Region** | **Cluster size** | **Peak t-value** | **Peak p-value (FDR)** | **MNI coordinates (x, y, z)** | **Hedges’ g [Bootstrap 95%CI]** | **BF10** |
|---|---|---|---|---|---|---|
| **Vis. > Aph.** |  |  |  |  |  |  |
| Left aPFC | 52 | 4.68 | 0.000 | -36, 54, 4 | 1.53 [0.87, 2.53] | 433.22 |
| Left PoCG | 49 | 3.73 | 0.005 | -64, -14, 22 | 1.21 [0.55, 2.22] | 41.42 |
| **Aph. > Vis.** |  |  |  |  |  |  |
| Left PHG/EC | 264 | 6.83 | 0.000 | -25, -19, -23 | -2.23 [-3.27, -1.60] | 132680.97 |
| Left ACC | 44 | 3.61 | 0.005 | -4, 18, 23 | -1.18 [-2.03, -0.57] | 31.67 |
| Right RSC | 294 | 3.52 | 0.007 | 17, -53, 6 | -1.15 [-1.88, -0.57] | 26.20 |
| Right EC/IFG | 145 | 4.49 | 0.000 | 32, -6, -38 | -1.46 [-2.30, -0.86] | 266.52 |
| Right PCC | 110 | 4.41 | 0.001 | 5, -18, 29 | -1.44 [-2.17, -0.90] | 214.95 |

**Supplementary Table S6.** Levene’s test for equality of variances. Group differences of variance were examined across all main variables identified using Levene’s test.

| Variables | Vis SD | Aph SD | p value (uncorrected) |
|---|---|---|---|
| ***Tractometry*** |  |  |  |
| right UF | 0.028 | 0.041 | 0.088 |
| left UF | 0.032 | 0.045 | 0.191 |
| interparietal corpus callosum | 0.027 | 0.037 | 0.293 |
| right dorsal cingulum | 0.036 | 0.037 | 0.793 |
| left dorsal cingulum | 0.038 | 0.035 | 0.717 |
| ***Graph theory*** |  |  |  |
| Strength left FEF | 6.053 | 7.975 | 0.535 |
| Clustering coefficient left anterior insula | 0.002 | 0.002 | 0.761 |
| Clustering coefficient left dlPFC | 0.004 | 0.004 | 0.919 |
| Participation coefficient right somatomotor | 0.049 | 0.086 | 0.219 |
| Participation coefficient right dmPFC 1 | 0.049 | 0.033 | 0.136 |
| Participation coefficient right dmPFC 2 | 0.029 | 0.018 | 0.445 |
| ***Thickness*** |  |  |  |
| Left aPFC | 0.097 | 0.117 | 0.478 |
| Left PoCG | 0.117 | 0.142 | 0.535 |
| Left PHG/EC | 0.36 | 0.401 | 0.463 |
| Left ACC | 0.176 | 0.189 | 0.892 |
| Right RSC | 0.299 | 0.231 | 0.301 |
| Right EC/IFG | 0.36 | 0.401 | 0.463 |
| Right PCC | 0.101 | 0.136 | 0.335 |

### Multiple regression model

We used a two-step modelling approach. First, LASSO-penalized logistic regression was applied separately within each metric domain to identify the most informative predictors. Second, the variables retained by LASSO were entered into Firth’s penalized logistic regression to obtain bias-reduced parameter estimates and odds ratios.

For the tractometry model, LASSO logistic regression retained the left dorsal cingulum, right uncinate fasciculus, and posterior interparietal corpus callosum at λ se. In the Firth model, higher FA in the right uncinate fasciculus was marginally associated with lower odds of aphantasia (beta = −1.00; OR = 0.36, 95% CI = 0.07-1.00; p = 0.051), as was higher FA in the posterior interparietal corpus callosum (beta = −1.70; OR = 0.18, 95% CI = 0.02-0.61; p = 0.003). Higher FA in the left dorsal cingulum was significantly associated with increased odds of aphantasia (beta = 1.79; OR = 6.03, 95% CI = 1.79-51.40; p < 0.001). These multivariate results were broadly consistent with the tract-level univariate findings. Cross-validation area under the curve (CV-AUC) for this model was 0.87 (95% CI 0.82–0.91).

For the cortical thickness model, we selected the left anterior prefrontal cortex (aPFC) and right retrosplenial cortex (RSC) clusters, which were the largest clusters showing negative and positive effects, respectively, to avoid complete separation in the logistic regression model. LASSO logistic regression retained both clusters. In the final Firth model, reduced left aPFC thickness (beta = −1.81; OR = 0.16, 95% CI = 0.03–0.48; p < 0.001) and greater right RSC thickness (beta = 1.52; OR = 4.58, 95% CI = 1.54–27.33; p = 0.004) were associated with higher odds of aphantasia. The area under the curve (AUC) for the cortical thickness model was 0.93. In addition, we examined the single-variable effect of the left parahippocampal gyrus/entorhinal cortex (PHG/EC) cluster, which showed the largest effect size. Greater left PHG/EC thickness contributed to group discrimination (beta = 2.99; OR = 19.90, 95% CI = 4.27–313.68; p < 0.001). The CV-AUC for this model was 0.90 (0.87–0.92).

For the graph-theoretical metrics model, given concerns that including all candidate predictors would result in an overly complex model relative to the sample size, we excluded the participation coefficient in the right dorsomedial prefrontal cortex (dmPFC, region 3) and the somatomotor cortex before applying LASSO, based on overlap among regions and their relatively lower theoretical relevance to aphantasia. LASSO regression retained four predictors spanning three network measures. In the final Firth model, a lower clustering coefficient in the left insula (β = −2.41; OR = 0.09, 95% CI = 0.001–0.47; p = 0.002), a lower clustering coefficient in the left prefrontal cortex (β = −0.89; OR = 0.41, 95% CI = 0.007–2.17; p = 0.336), and lower node strength in the left frontal eye field (FEF) (β = −1.32; OR = 0.27, 95% CI = 0.029–0.84; p = 0.022) were associated with higher odds of aphantasia. A higher participation coefficient in the right dmPFC was also associated with higher odds of aphantasia (β = 1.81; OR = 6.14, 95% CI = 1.29-216.11; p = 0.021), consistent with the corresponding univariate result. The CV-AUC for the graph-theoretical metrics model was 0.93 (0.87–0.96).

## Methods

### Participants

Eighteen individuals with self-reported congenital aphantasia (mean ± SD age, 31.79 ± 12.11 years; 11 female) and 18 typical visualizers (32.29 ± 9.53 years; 15 female) were recruited. Ten aphantasic participants and five visualizers had also participated in a previous 7 Tesla fMRI study of mental imagery (Liu, Zhan, et al., 2025). The sample size of 18 per group reflects the recruitment constraints inherent to congenital aphantasia, which affects an estimated 2–4% of the population. This is comparable to landmark structural MRI studies of analogous congenital conditions (Thomas et al., 2009). All participants completed the Vividness of Visual Imagery Questionnaire (VVIQ; Marks, 1973), which comprises 16 items rated from 1 (“no image at all”) to 5 (“perfectly clear and as vivid as normal vision”), for a total score ranging from 16 to 80. All aphantasic participants had VVIQ scores below 22 (mean ± SD, 17.06 ± 2.10; core aphantasia, 13 out of 18), and a lifelong absence of voluntary visual imagery was confirmed by a structured interview conducted by J.L. No widely accepted objective diagnostic marker of congenital aphantasia is currently available (Bouyer et al., 2025). All visualizers had VVIQ scores above 50 (63.67 ± 9.54). All participants were right-handed, had normal or corrected-to-normal vision and hearing, and reported no history of neurological or psychiatric disorder. The two groups were matched for age, sex, and years of education. The study was approved by the institutional review board of INSERM (protocol C13-41), and all participants provided written informed consent in accordance with the Declaration of Helsinki.

### Image acquisition

All MRI data were acquired at the Centre de NeuroImagerie de Recherche (CENIR), Paris Brain Institute, on a Siemens 3T MAGNETOM Prisma scanner equipped with a 64-channel head coil. Diffusion-weighted imaging (DWI) data were acquired across four sequences (398 volumes total), using two phase-encoding directions (anterior-to-posterior, AP; posterior-to-anterior, PA), each with 98-99 uniformly distributed diffusion directions at two b values (b = 1,500 and 3,000 s/mm²). In addition, 28 non-diffusion-weighted (b = 0) volumes were acquired. High-resolution T1-weighted anatomical images were acquired in the same session (TR = 2,400 ms, TE = 2.22 ms, flip angle = 9°).

### DWI and T1-weighted data processing

#### Grey matter/white matter segmentation

Anatomical T1-weighted images were preprocessed using the DeepPrep 25.1.0 pipeline (Ren et al., 2025), which combines deep-learning-based and established neuroimaging tools for robust tissue segmentation. Processing included motion correction using FreeSurfer (v7.2.0; RRID: SCR_001847), bias-field correction using SimpleITK (v2.3.0), skull stripping and tissue segmentation using FastSurferCNN from FastSurfer (v1.1.0), and cortical surface reconstruction using FastCSR (v1.0.0). The preprocessed T1-weighted image served as the anatomical reference for subsequent diffusion-weighted analyses.

#### Diffusion-weighted image preprocessing

Diffusion-weighted images were preprocessed using QSIPrep v1.0.2 (Cieslak et al., 2021) with default parameters. Sequences with the same phase-encoding polarity were merged before preprocessing. The pipeline included MP-PCA denoising, Gibbs-ringing removal, intensity normalization across b = 0 images, N4 bias-field correction, and correction for head motion, eddy currents, and susceptibility-induced distortion using FSL’s eddy and TOPUP, with outlier replacement enabled. Final interpolation used Jacobian modulation, and the data were resampled to each participant’s native AC-PC space at 1.25 mm isotropic resolution.

#### Fiber reconstruction

Preprocessed DWI data were reconstructed using QSIRecon v1.0.1 (Cieslak et al., 2021) with the *mrtrix_multishell_msmt_ACT-hsvs* workflow. Brain masks from *antsBrainExtraction* were used throughout reconstruction. Multi-tissue response functions were estimated using the Dhollander algorithm, and white-matter fiber orientation distributions (FODs) were computed using multi-tissue constrained spherical deconvolution and intensity-normalized with *mtnormalize*. These FODs were used for tractography, whereas anatomical constraints were derived from T1-weighted hybrid surface-volume segmentation (hsvs) to generate five-tissue-type images. Whole-brain tractography was then performed using anatomically constrained tractography (ACT), with streamlines constrained to initiate and terminate at the grey-white matter interface. SIFT2 was applied to estimate per-streamline weights, which were retained for subsequent bundle segmentation and connectivity analyses.

#### Bundle segmentation

White-matter fascicles of interest were identified using pyAFQ (Kruper et al., 2021) within QSIRecon, following the *mrtrix_multishell_msmt_pyafq_tractometry* workflow with ACT. In addition to the standard major fiber bundles included in pyAFQ, several custom tracts were defined for hypothesis-driven analyses using region-of-interest (ROI) definitions from prior studies. These included superior longitudinal fasciculus subdivisions (Thiebaut de Schotten et al., 2011) (SLF I, II, III) and ventral cingulum pathways containing posterior cingulate and medial temporal projections (Warrington et al., 2020). All ROIs were transformed into each participant’s native AC-PC space. Association fibers were constrained to remain within the hemisphere, and bundle refinement was performed using iterative outlier removal and streamline clustering.

### Tractometry

For each segmented bundle, tract profiles were extracted by resampling streamlines to 100 equidistant nodes using the pyAFQ workflow. Node-wise diffusion and microstructural metrics were then computed as weighted averages across streamlines, including fractional anisotropy (FA), mean diffusivity (MD), radial diffusivity (RD), axial diffusivity (AD), and neurite orientation dispersion and density imaging (NODDI) parameters. FA is sensitive to overall fiber organization but does not distinguish between true microstructural differences and geometric effects such as crossing, fanning, or bending fibers (Sotiropoulos & Zalesky, 2019). To aid interpretation, we therefore also quantified NODDI-derived neurite density index (NDI) and orientation dispersion index (ODI), which provide more specific estimates of tissue microstructure (H. Zhang et al., 2012). NODDI was fitted using the AMICO implementation in QSIRecon, with parallel diffusivity fixed at 1.7 × 10^-3 mm²/s and isotropic diffusivity at 3.0 × 10^-3 mm²/s. NDI and ODI values were computed at each node along the tract profiles. Prior to group-level statistical analysis, tract profiles were smoothed along the tract using a 3-node full-width at half maximum (FWHM) kernel.

### fROI fiber tracking

Four ROIs from a core mental imagery network were extracted from a previous 7 Tesla fMRI study of aphantasia (Liu, Zhan, et al., 2025): the fusiform imagery node (FIN), left anterior prefrontal cortex (aPFC), left orbitofrontal cortex (OFC), and right anterior intraparietal sulcus (aIPS). In aphantasic individuals, these ROIs showed altered functional connectivity or activity, motivating the analysis of their white-matter connections. Group-level ROIs were transformed into each participant’s native AC-PC space. To avoid overestimation of ROI size, volumetric fROIs were projected onto the cortical surface using FreeSurfer (RRID:SCR_001847) and mapped to the grey-white matter interface (Meisler et al., 2024). The streamline search space was defined as the intersection between the projected fROIs and the grey-white matter interface mask. Streamlines intersecting this region were identified using MRtrix3 (mrcalc) with the default radial endpoint search radius of 2 mm. For each ROI, streamlines were extracted and quantified in native space using two indices: the proportion of streamlines assigned to each anatomically defined bundle, and the number of streamlines normalized by ROI volume. Bundles of interest were defined according to the anatomical location of each ROI. For the FIN, these included the long segment of the arcuate fasciculus (AF), posterior arcuate fasciculus (pAF), inferior fronto-occipital fasciculus (IFOF), inferior longitudinal fasciculus (ILF), and vertical occipital fasciculus (VOF). For the left aPFC and left OFC, the bundles of interest were the IFOF and uncinate fasciculus (UF). For the right aIPS, the bundles of interest were the pAF and SLF I, II, and III. Group differences were assessed using both normalized streamline counts and bundle-wise streamline proportions.

### Graph-theoretic network analysis

Cortical parcellation was performed using the Schaefer 400-parcel, 17-network atlas (Schaefer et al., 2018), registered from template space to individual DWI space via T1-weighted normalization. Structural connectivity between parcel pairs was estimated by intersecting the endpoints of SIFT2-weighted streamlines with parcel ROIs. A streamline was assigned to a parcel pair if both endpoints fell within a 2 mm radius sphere centered on the corresponding parcel centroids. Connection strength was defined as the sum of SIFT2 weights across all streamlines linking a parcel pair, normalized by parcel volume to account for region-size differences. Symmetric adjacency matrices were generated for each participant and thresholded using consensus thresholding (de Reus & van den Heuvel, 2013), such that connections absent in at least 50% of participants in either group were set to zero. Graph-theoretic metrics were computed on weighted, undirected networks using the Brain Connectivity Toolbox (Rubinov & Sporns, 2010) in MATLAB R2024b (RRID: SCR_001622). The metrics examined were node strength, clustering coefficient, global efficiency, betweenness centrality, and participation coefficient. To assess robustness across parcellation schemes, analyses were repeated using the Schaefer 100-parcel and Gordon 333-parcel functional atlases.

### Surface extraction and surface-based morphometry

Surface-based morphometry was conducted using the Computational Anatomy Toolbox (CAT12) implemented in SPM12 (Gaser et al., 2024). Cortical thickness was estimated using CAT12’s default projection-based method, which computes the distance between the inner cortical surface (grey-white matter boundary) and the outer cortical surface (grey matter-cerebrospinal fluid boundary). Individual cortical thickness maps were smoothed with a 12 mm FWHM Gaussian kernel and resampled to the 32k surface before statistical analysis. Whole-brain statistical analysis was performed in CAT12. The resulting t maps were then used as input for Neurosynth (RRID:SCR_006798) decoding in BrainStat to compute Pearson correlations with meta-analytic activation maps. The 20 strongest term correlations were retained for visualization.

To examine the visual cortex specifically, we used two atlas-based approaches. Grey matter volume was extracted from occipital regions using the Neuromorphometrics atlas (RRID: SCR_005656) as implemented in CAT12, and cortical thickness was extracted from early visual regions using the HCP-MMP Glasser atlas. Grey matter volume measures were normalized by total intracranial volume to account for individual differences in head size.

### Quantification of NODDI in grey matter and white-matter tracts

To complement cortical thickness analysis with microstructurally interpretable measures, NODDI metrics were also quantified in cortical grey matter. NODDI was fit using the AMICO implementation described above, and volumetric NDI and ODI maps were sampled onto the cortical surface uniformly between the pial and white matter boundaries using Nilearn’s *vol_to_surf* function, then resampled to the fsLR-32k surface. The NODDI model has been validated against histological measures in both grey and white matter, and ODI has been shown to be sensitive to microstructural changes including demyelination in post-mortem tissue.

### Logistic regression

To identify parsimonious combinations of structural features associated with group membership while accounting for shared variance across measures, we used a two-step modelling approach. First, LASSO-penalized logistic regression (*glmnet* package in R) was applied separately within each metric domain: tractometry FA, cortical thickness, and graph-theoretic measures. Predictors comprised mean FA across significant tract nodes, cortical thickness values, and graph metrics identified in prior analyses. Separate domain-specific models were used to limit the predictor-to-sample ratio given the modest sample size (N = 36). Model selection was performed using 5-fold cross-validation with approximately balanced class distributions, and the regularization parameter was chosen using the 1-standard-error rule (λ se). In a second step, variables retained by LASSO were entered into Firth’s penalized logistic regression (Firth, 1993) to obtain bias-reduced parameter estimates and odds ratios, thereby reducing small-sample bias and separation problems. Variables that caused non-convergence were excluded using a p-value-based stepwise procedure. To assess out-of-sample discrimination and guard against overfitting, each domain model was evaluated with repeated stratified cross-validation (1000 repetitions of stratified 5-fold cross-validation), refitting the model on each training partition and quantifying discrimination from out-of-fold predictions.

### Statistical analysis

#### Bayes factors

To quantify evidence for and against group differences, we computed Bayes factors (BFs) in JASP (https://jasp-stats.org/; RRID: SCR_015823) using Bayesian t tests and Mann-Whitney U tests with default priors on effect size. BF10 > 3 was taken as moderate evidence for a group difference, whereas BF01 > 3 (equivalently, BF10 < 1/3) was taken as moderate evidence for the absence of a group difference (Keysers et al., 2020).

#### Effect size

For significant findings, effect sizes were calculated from the mean values within each significant region. Hedges’ g was used for FA, cortical thickness, and NODDI metrics as a bias-corrected standardized effect size to reduce small-sample bias. Cliff’s delta was used for graph-theoretic metrics as a nonparametric measure of effect size. 95% confidence intervals for all effect sizes were estimated using 10,000 bootstrap resamples.

#### Tractometry

Group differences in tract profiles were assessed using one-dimensional threshold-free cluster enhancement (TFCE; E = 0.5, H = 2) along tract nodes, implemented in PALM with 10,000 permutations. TFCE identifies spatially extended effects without requiring an arbitrary cluster-forming threshold and assigns a family-wise-error-corrected p value to each node. To reduce spurious findings, only clusters comprising at least 3 consecutive significant nodes were reported. Statistical significance was defined as p < 0.05, FWE-corrected.

#### Graph-theoretic network analysis

Group differences in node-level graph-theoretic metrics were assessed using the non-parametric Brunner-Munzel test, with false-discovery-rate correction applied across all nodes within each metric. Furthermore, to assess whether the findings on node-level networks reflected topology-specific differences or more general properties of the weighted structural connectome, we performed a null-network sensitivity analysis using the Brain Connectivity Toolbox (Rubinov & Sporns, 2010). This analysis focused on the weighted clustering coefficient, because clustering measures are sensitive to local network density, degree distribution, and weight distribution. For each participant, 100 null networks were generated from the empirical weighted undirected connectivity matrix using the *“null_model_und_sign”* function. This function randomizes the network while preserving the degree, weight, and strength distributions of the original matrix. In the present analysis, the number of binary rewiring swaps per edge was set to 5, and the frequency of weight sorting during weighted randomization was set to 0.1. For each participant and region, the empirical clustering coefficient was normalized by the mean clustering coefficient obtained from the corresponding null-network distribution. Specifically, the null-normalized value was calculated as the ratio of the empirical value to the mean of the null values. The resulting null-normalized clustering coefficient values were compared between groups using the same statistical procedure as in the original node-level network analysis.

#### fROI fiber tracking

Group differences in filtered tracts within each fROI were assessed using the Mann-Whitney test, because the data included a substantial number of zero values and did not satisfy the assumptions of parametric testing.

#### Surface-based morphometry

Statistical analyses of cortical thickness were performed in BrainStat (Larivière et al., 2023) using a linear model with a one-sided test and false discovery rate (FDR) correction. To identify spatially coherent clusters of significant vertices, hierarchical density-based spatial clustering of applications with noise (HDBSCAN) was applied to the x-, y-, and z-coordinates of significant vertices, grouping those that were spatially adjacent. Only clusters containing at least 30 vertices were retained for interpretation, and Desikan killiany atlas was used to identify the anatomical region of surviving clusters.

#### Grey matter NODDI

Cortical grey matter NODDI metrics were compared between groups using random field theory as implemented in BrainStat (Larivière et al., 2023). Cluster-level inference used a cluster-forming threshold of p < 0.05 and a cluster-significance threshold of p < 0.05, FWE-corrected.

#### Levene’s test for equality of variances

To assess the heterogeneity of aphantasia profile in the current samples, Brown–Forsythe alternative of the Levene’s test was applied separately within each metric.

